# multiDGD: A versatile deep generative model for multi-omics data

**DOI:** 10.1101/2023.08.23.554420

**Authors:** Viktoria Schuster, Emma Dann, Anders Krogh, Sarah A. Teichmann

## Abstract

Recent technological advancements in single-cell genomics have enabled joint profiling of gene expression and alternative modalities at unprecedented scale. Consequently, the complexity of multi-omics data sets is increasing massively. Existing models for multi-modal data are typically limited in functionality or scalability, making data integration and downstream analysis cumbersome. We present multiDGD, a scalable deep generative model providing a probabilistic framework to learn shared representations of transcriptome and chromatin accessibility. It shows outstanding performance on data reconstruction without feature selection. We demonstrate on several data sets from human and mouse that multiDGD learns well-clustered joint representations. We further find that probabilistic modelling of sample covatiates enables post-hoc data integration without the need for fine-tuning. Additionally, we show that multiDGD can detect statistical associations between genes and regulatory regions conditioned on the learned representations. multiDGD is available as an scverse-compatible package (https://github.com/Center-for-Health-Data-Science/multiDGD).

## 1 Introduction

Single-cell genomics methods have become the main technology to study cellular heterogeneity and dynamics within tissues. They also enable the measurement of multiple molecular features within individual cells, pairing measurements of the transcriptome with epigenome, proteome or genome profiling. These paired multi-modal measurements can be used for deeper characterization of cell states, differentiation processes or genotype-to-phenotype relationships [1].

Analysis of paired single-cell multi-omics data typically requires joint dimensionality reduction on multiple molecular measurements to identify cell-cell similarities, cell states and patterns of co-variation between genomic features (also known as vertical integration [2]). Several statistical models have been proposed for this task, mostly based on factor analysis [3–5] or cell-cell similarity embeddings [6, 7]. Recently, approaches have been proposed to additionally integrate paired data from measurements of individual modalities (i.e. mosaic integration) [8–11]. However, existing methods have primarily been applied to relatively small data sets, while increasing availability of multimodal data now requires models that can handle tens of thousands of cells from multiple experiments, with the ability to account for technical differences between samples [12]. Additionally, methods for vertical integration struggle with imbalance in the dimensionality of feature spaces, especially in the joint analysis of gene expression and chromatin accessibility over hundreds of thousands of genomic regions [2]. Importantly, as the field and its methodologies are still developing, existing analytical approaches predominantly focus on dimensionality reduction for the purpose of cell clustering, with notably little emphasis on identifying relationships between molecular features [1]. This is especially relevant for joint analysis of epigenomic and transcriptomic profiles to associate regulatory regions to changes in gene expression.

In order to alleviate the problems encountered with large data sets, generative models have been applied to both uni-modal [13–18] and multi-modal [8, 11, 19–21] data. Deep generative models are powerful machine learning techniques that aim to learn the underlying function of how data is generated. This is of special interest for unsupervised analysis of single-cell data, where the goal is to interpret patterns of variation in high-dimensional and noisy data [22]. The predominant type of generative model applied in this field is the Variational Autoencoder (VAE) [23]: models tailored for scRNA-seq data [13–16] enable integration of large and complex data sets at lower computational cost [24] and have been successfully applied to the analysis of cells across human tissues and in large cohorts [25–27]. These models do however come with some limitations [18], which are continuously being addressed by a large community. With the current state of model design, it is for example not trivial to integrate new samples from different batches after training as covariates are modelled via one-hot encoding. scArches [28] is a tool introduced to solve this problem post-hoc, but the proposed fine-tuning is rather a band-aid than a solution to the underlying problem.

While the number of generative models available is vast for scRNA-seq single-cell data [13–18, 21], the application to multi-modal single-cell data has just begun. Existing models often employ simplistic architectures, priors for the generative distribution and encoding of confounding covariates such as batch effects [8, 11, 19, 20]. The results are under-performing models which hold the noise in the data accountable[29]. Generative modelling can provide much more than a joint integration. As emergent properties, deep generative models can capture underlying relationships between variables and dynamics in high-dimensional data. Straightforward examples of this would be feature interactions and cell state transitions. These properties can be learned without explicit modelling. However, these promising applications of generative models are still under-explored, as many models focus only on a fraction of the actual feature space.

In this work, we propose a new generative model, multiDGD, which aims to provide a basis for improved data integration and analysis of feature interactions. The model is an extension of the Deep Generative Decoder (DGD) [18] for single-cell multi-omics data. Unlike VAE-based models, it uses no encoder to infer latent representations but rather learns them directly as trainable parameters, and employs a Gaussian Mixture Model (GMM) as a more complex and powerful distribution over latent space. This introduces several advantages. Firstly, an encoder limits the flexibility and quality of representations. A decoder alone can better recover representations close to the optimum and reduces the number of parameters in the model [30]. Secondly, our GMM as a distribution over latent space increases the ability of the latent distribution to capture clusters in comparison to the standard Gaussian used in applied VAEs. Another strength of the DGD is its data efficiency. As presented in [30], the encoder requires more data to be well defined than the decoder. Removing the encoder makes the model applicable to not only large but also small data sets. This also translates to the number of features that can be modelled, and makes the DGD amenable to model genome-wide chromatin accessibility data where feature selection is problematic and may not be desirable.

We demonstrate on real world applications that the DGD can learn meaningful representations of complex multi-modal data, with improved performance for dimensionality reduction, cross-modality prediction, and modelling of unseen batches without the need for fine tuning. Furthermore, we provide a proof-of-concept that multiDGD can be used to predict regulatory associations between genes and peaks based on *in silico* perturbation.

## 2 Results

### 2.1 The model

multiDGD is a generative model of transcriptomics and chromatin accessibility data. It consists of a decoder mapping shared representations of both modalities to data space, and learned distributions defining latent space. Fig. 1 shows a schematic of multiDGD with its training and inference processes.

**Fig. 1.**
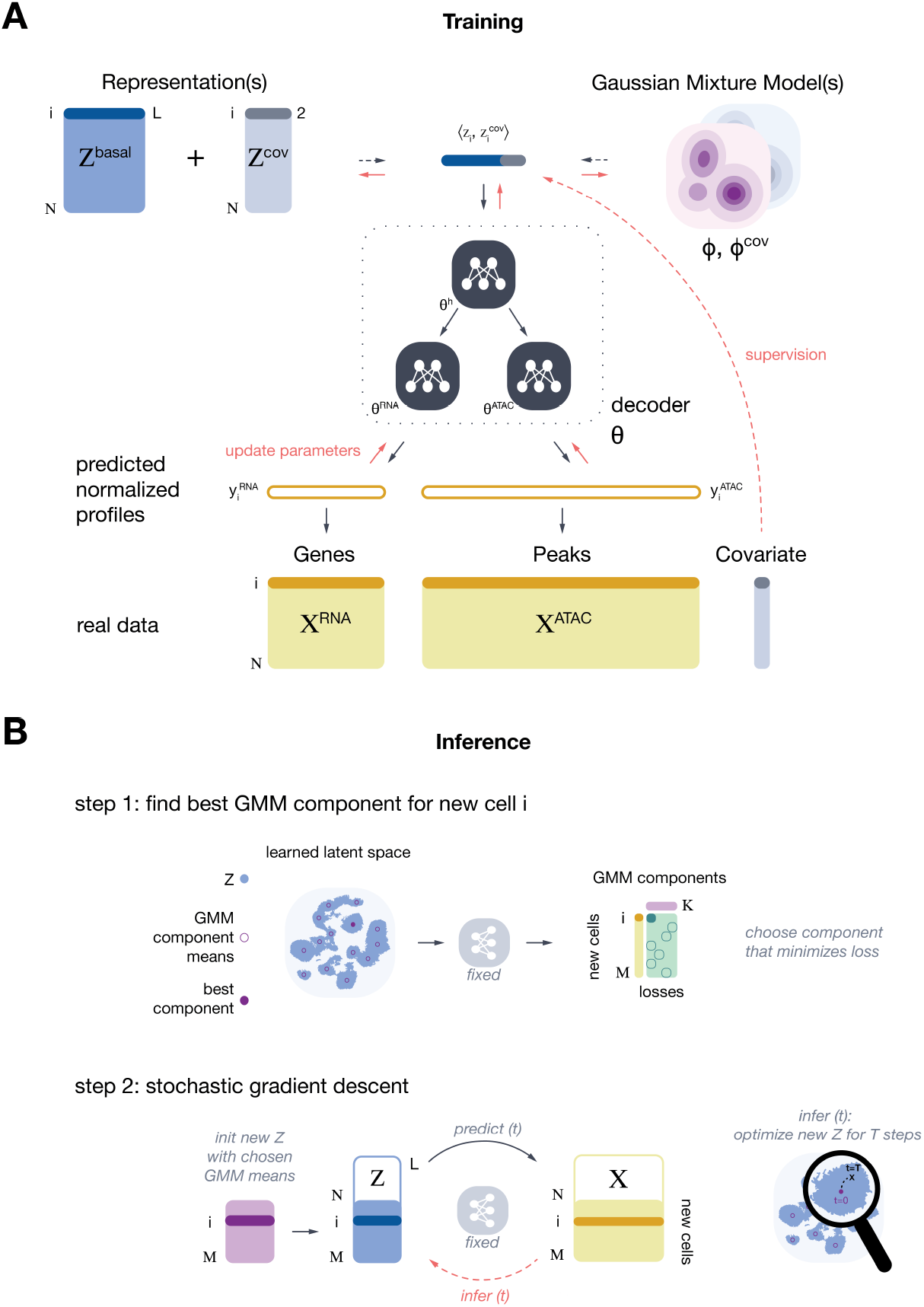
Schematic and graphical model of multiDGD. **A)** Schematic of model architecture and generative process. Representations *Z* present the input to the decoder. They are distributed in latent space according to a Gaussian mixture distribution parameterized by *ϕ*. While *Z*^*basal*^ and *ϕ* are required, *Z*^*cov*^ and *ϕ*^*cov*^ are optional and depend on the presence of covariates to be modelled. For each data point (cell) *i ∈ N*, there exists a latent vector of length L, plus 2 dimensions for each covariate modelled. The input is transformed into modality-specific predicted normalized mean counts *y* through the hierarchical decoder *θ*. These outputs are then scaled with sample-wise count depths to predict the density of both RNA and ATAC data. Orange arrows depict the backpropagation and updating of parameters during training. **B)** Schematic of the two-step process of inferring new representations. In step 1, the best mode of *ϕ* is selected for each new sample *i∈ M* as the argmax of the probability densities of each component *k∈ K*. In step 2, each initial representation is optimized for a specified number of steps *T* with fixed model parameters.

The inputs to the decoder are the low-dimensional representations *Z* of data *X*. Instead of providing them through an encoder (as in the Variational Autoencoder [23]), they are learned directly as trainable parameters [30]. One of the extensions to the model is the inclusion of disentangled representations. As a result, we can model the ‘molecular’ representation of cells *Z*^*basal*^ separately from technical batch effects and sample covariates (*Z*^*cov*^). Representations *Z* are concatenations of the two. Parameters *ϕ* and *ϕ*^*cov*^ represent the parameterized distributions over latent space. They could be represented by any differentiable distribution, but we chose Gaussian Mixture Models (GMMs), as we see them as a natural choice for data containing sub-populations and can provide unsupervised clustering.

Data is generated by feeding latent representations *Z* to the decoder. For every *i*th sample of *N* data samples (cells), there exists a corresponding representation *z*_*i*_. The decoder consists of three blocks: the shared neural network (NN) *θ*^*h*^, and the two modality-specific NNs *θ*^*RNA*^ and *θ*^*AT AC*^. The modality-specific networks predict fractions of the total counts per cell and modality, *y*_*ij*_. These are then converted into predicted means of Negative Binomial distributions modelling counts by multiplying with the total count *s*_*i*_. The training objective is given by the joint probability *p*(*X, Z, θ, ϕ*) [18], which is maximized using Maximum a Posteriori estimation [18]. Both the model and the inference process are explained in more detail in the Methods section and Supplementary Figure 6.

### 2.2 Improved performance on data reconstruction and clustering

We first evaluate this model’s performance compared to VAE-based alternatives. Since MultiVI [8], a VAE-based multi-omics generative model, presents the only one which can integrate data from different batches and impute missing data, it is the main focus of our benchmark. Where applicable, we included performances of Cobolt [11] and scMM [20]. We compare performances for three different data sets of paired scRNA-seq and scATAC-seq, derived from human bone marrow [31], human brain [32] and mouse gastrulation [33] multiomics data (see Methods). Comparing the data reconstruction performances on a held out test set, we found that multiDGD consistently outperforms MultiVI on all tested data sets (as well as scMM on the brain data) (Figure 2A-B). The improvement is especially large for the reconstruction of ATAC features.

**Fig. 2.**
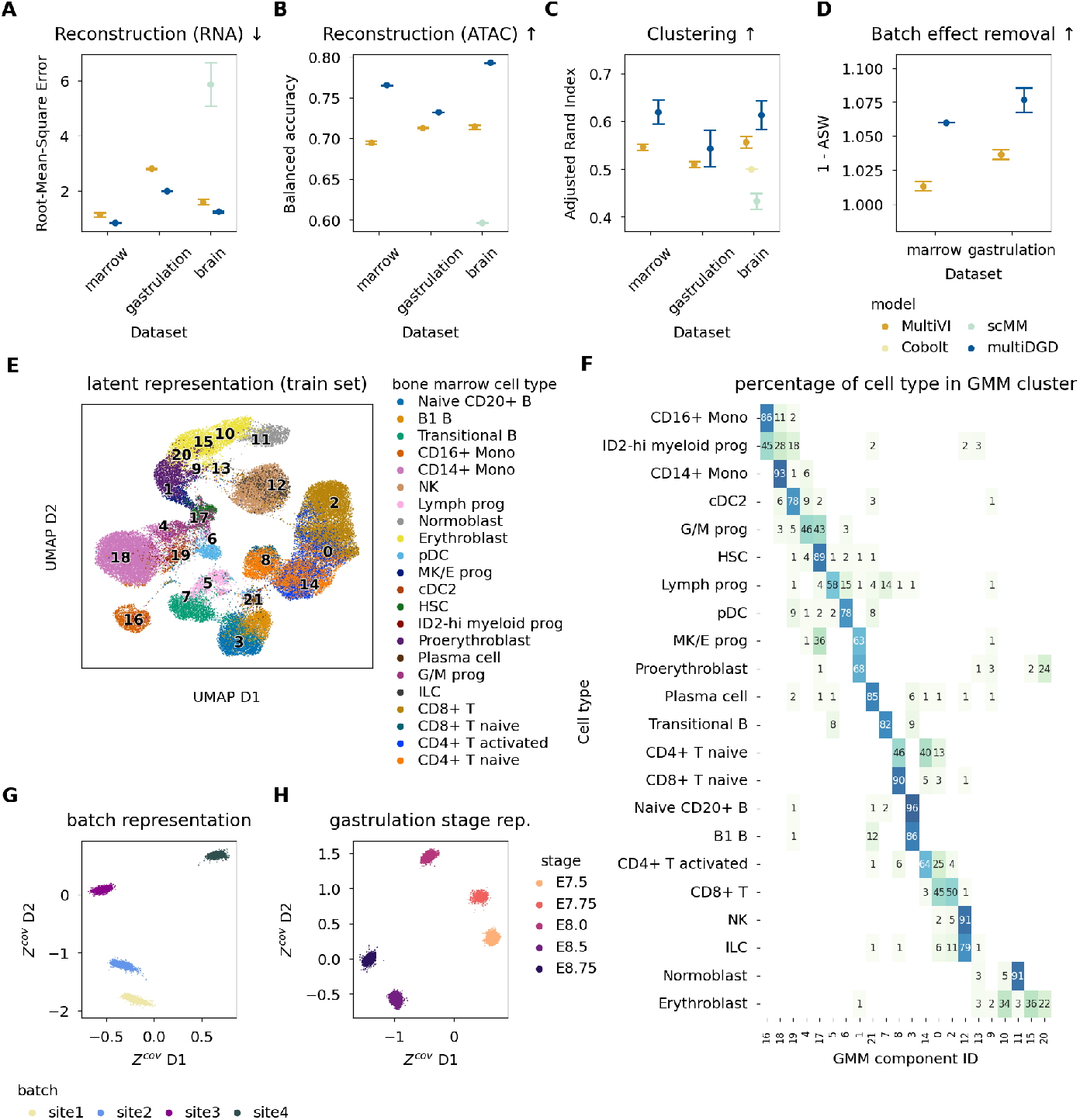
High performance on reconstruction, clustering and the addition of probabilistic models for covariates. **A-D)** Performance evaluations were done on the three data sets marrow, gastrulation and brain, and three different random seeds for all models. Cobolt [11] and scMM [20] were included where applicable. Cobolt does not provide data reconstruction/imputation, and both Cobolt and scMM do not include batch correction. MultiVI [8] was used as the main comparison. **A)** Reconstruction performance on the test RNA data measured by RMSE. Lower is better. **B)** Comparison of the reconstruction performance on the test set ATAC data as the balanced accuracy of binarized predictions. Higher is better. **C)** Clustering performance of the train latent spaces. The metric is the ARI based on clustering derived from the GMM for multiDGD and Leiden clustering for MultiVI. The Leiden algorithm is adjusted so that the number of clusters in both models’ latent spaces are comparable. More in Methods. Higher is better. **D)** Batch effect removal of marrow and gastrulation data was calculated as 1 *− ASW* . Brain was not included as the data annotation contained no batch information. **E-G)** Latent space visualizations and clustering evaluation of the bone marrow latent spaces. **E)** UMAP visualization of the multiDGD latent representation. It is colored by annotated cell types. GMM component means are indicated by black numbers projected onto their transformed coordinates. **F)** Heatmap of the clustering of representation samples grouped by annotated cell types. Numbers inside the heatmap indicate the percentage of samples in a cell type assigned to a given component. **G)** Covariate representations colored by Site. The legend is the same as in H. **H)** Covariate representations of the gastrulation stage from mouse gastrulation data.

We next evaluated the latent spaces learned by the models in terms of clustering of cell types and batch effect removal. MultiVI’s shared embeddings of transcriptional and chromatin features are commonly used as input for the Leiden [34] clustering algorithm. The DGD intrinsically performs clustering with the Gaussian Mixture Model as the latent distribution (details in Methods sections 4.2.3,4.2.11). Measuring clustering performances with the Adjusted Rand Index (ARI) (Figure 2C), we see a notable variance in performance with random seeds for model initialization, more so for multiDGD than for MultiVI. However, the GMM components of multiDGD still learn latent representations whose clustering generally aligns better with the annotated cell type labels (compared to MultiVI, Cobolt and scMM). Figures 2E-F visualize the intrinsic clustering on the example of the human bone marrow benchmark data (remaining data sets are shown in Supplementary Figures 8 and 13). For some lineages, individual cell types are harder to separate than others. One of the difficult cell types in general are T cells [35]. Unlike many of the other cell types, the four subtypes do not cluster according to the four T-cell-associated GMM components. This might be due to limitations of RNA expression as a discriminant of T cell subtypes. Nevertheless, multiDGD clusters associated with T cells contain less spillover than Leiden clusters derived from MultiVI latent representations (Supplementary Figures 9 and 10). We see a clear distinction between naive CD8 T cells in component 8 and activated CD8 T cells in components 0 and 2. We also observe that very small cell type clusters, such as progenitors, can be aggregated in a single component. Other components do not seem correlated with cell type identity and may rather model outliers (components 9, 13).

### 2.3 Probabilistic modelling of batch effects

Another important feature of generative models for single-cell data is the capability to alleviate batch effects. multiDGD leads to improved mixing between batches compared to MultiVI (Figure 2D and Supplementary Figure 8), although this is to be taken with a grain of salt as the average silhouette width may be skewed due to the different latent distributions. On our benchmark set, we see that the disentangled latent space results in clear separation of most cell types (Figure 2E and Supplementary Figure 8) and good mixture of the sites at which samples were processed (Supplementary Figure 11). The two-dimensional, separate representation for the batches derived from supervised training (Figure 2I) mirrors trends found in the general data distribution (see Supplementary Figure 12). These include site4 showing much more zero RNA counts than all other sites, which can explain why its cluster is distant from the others. In addition, we find that the covariate representation can capture biologically interpretable differences between samples. For example, when modelling the differences between embryos in the mouse gastrulation data set we see time-related structuring of the Gaussian components (Figure 2J). The early-to-late gastrulation phases from stage E7.5 to E8.0 [36] appear in chronological order, with stages E8.5 and E8.75 clearly separated. This distance makes sense as the differentiation of early organ progenitors is seen in stages E8.25 to E8.75 [36].

### 2.4 High performance on small data sets and many features

VAEs have clearly shown their usability and advantage when it comes to the speed at which they can model large data sets due to amortization, although this can come at the cost of posterior approximation [37]. Due to the missing encoder, the DGD is naturally suited for data sets with few samples and many features. Here, autoencoder-based models tend to overfit [30]. In this work, we briefly revisit this hypothesis by investigating the test performances of MultiVI and multiDGD trained on subsets of the human bone marrow set (Methods section 4.4.4). In order to put these into perspective, we compute average test loss ratios as the average test loss from the model trained on a subset over the test loss from the original model trained on the full set. Even though the variance in test loss ratios is much higher for multiDGD than for MultiVI (Figure 3), the average loss ratio of multiDGD stays stable for subsets larger than 1%, which corresponds to only 567 cells. MultiVI, on the other hand, performs worse with decreasing number of cells in the training data.

**Fig. 3.**
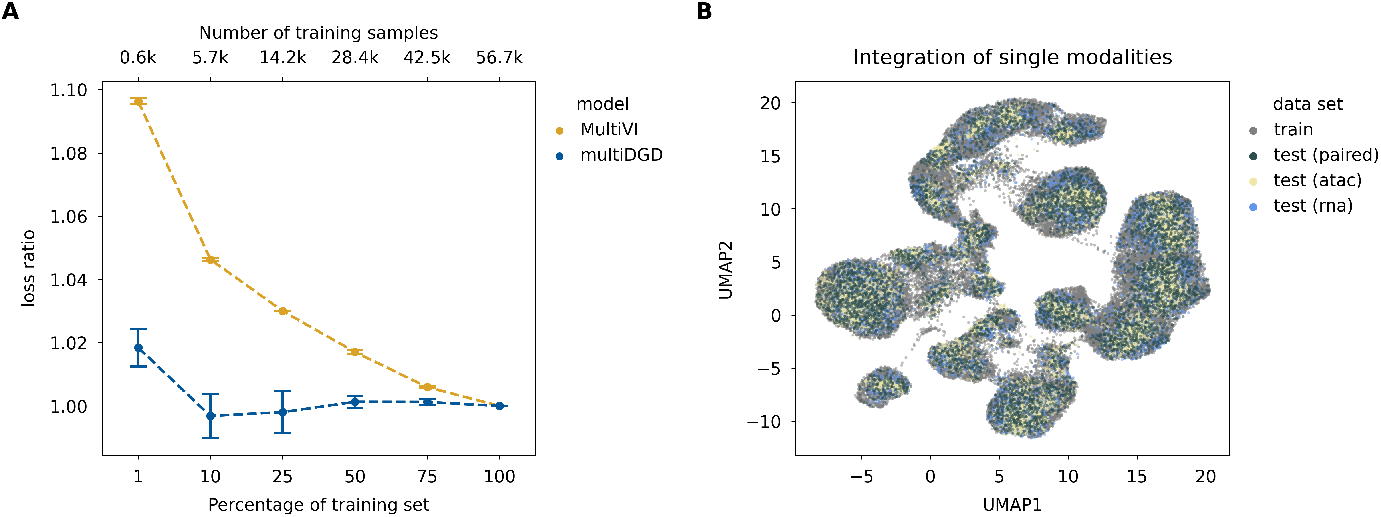
multiDGD is highly data efficient and can impute missing modalities. **A)** Performances on data efficiency are reported on both multiDGD and MultiVI. Data efficiency was evaluated by training the models designed for human bone marrow data on a variety of subsets. This was repeated for three different random seeds, resulting in the depicted error bars as the standard error of the mean. **B)** UMAP visualization of the full bone marrow latent space colored by sample type. We included the train and test splits, and the artificial uni-modal test representations ‘test (rna)’ and ‘test (atac)’.

This advantage of the DGD also carries over to data with many features. Typically, in single-cell analysis feature selection is applied before performing dimensionality reduction [38], both for scalability and to increase clustering performance [31]. While robust methods to select highly variable genes exist for scRNA-seq, there are no robust statistical methods for feature selection in scATAC-seq data sets. Here, accessibility is usually measured over hundreds of thousands of peaks, and several vertical integration methods suffer from this feature imbalance [2]. We compared multiDGD and MultiVI performance on data reconstruction in two scenarios on the mouse gastrulation data. The first one presents the previously presented models trained on data with feature selection (11792 genes, 69862 peaks), the second scenario presents models trained on all measured features (32285 genes, 192251 peaks). We compared performances on the shared set of features. While MultiVI lost performance for both modalities, multiDGD achieved nearly the same performance as before on ATAC data and even increased its performance on RNA data (Table 1).

**Table 1.** Feature efficiency on the mouse gastrulation set. Feature efficiency was investigated by training both models on two versions of the mouse gastrulation data. The first was the previously used set, where features had been selected for counts to be above zero for at least five percent of cells. The second version was the full data set. Performances were evaluated on the smaller set for comparability. We report the average performance metric with the standard error of the mean, RMSE for RNA data and balanced accuracy (BA) for ATAC data.

| Model | RMSE (RNA) | BA (ATAC) | Feature set |
| --- | --- | --- | --- |
| multiDGD | **1.7146 ± 0.0128** | **0.7382 ± 0.0007** | 5% |
| MultiVI | 2.3429 ± 0.0212 | 0.7121 ± 0.0003 | 5% |
| multiDGD | **1.6939 ± 0.0127** | **0.0.7397 ± 0.0007** | all |
| MultiVI | 2.4637 ± 0.0217 | 0.5197 ± 0.0001 | all |

**Table 2.** Summary of relevant symbols used to describe the DGD.

|  |  |
| --- | --- |
| $Z$ | representation |
| $X$ | data |
| $\hat{X}$ | predicted/ reconstructed data |
| mod | modality |
| $\theta$ | decoder parameters |
| $\phi$ | GMM parameters |
| $S$ | cell-specific scaling factor |
| $i \in N$ | single sample $i$ among $N$ total samples |
| $k \in K$ | component $k$ among $K$ components |
| $l$ | latent dimension |
| $c \in C$ | class $c$ in $C$ covariate classes |
| $\mu$ | GMM mean |
| $\Sigma$ | GMM covariance |
| $w, \pi$ | component coefficient, component weight |
| $\alpha$ | Dirichlet alpha |

### 2.5 Predicting and integrating unseen data

#### 2.5.1 Predicting missing modalities

We next evaluated the performance of multiDGD for prediction/imputation of missing data of one of the two modalities (RNA or ATAC). Predicting data modalities is a natural application of generative models for the case where existing uni-modal data is to be integrated with multi-modal data. In order to assess multiDGD’s predictive capability, we test its performance on the heldout test set given only one modality. Imputations are achieved by optimizing the partial likelihood of the available data described in Methods section 4.2.10. Fig. 3B shows that representations inferred from either the original paired samples or the artificial uni-modal samples are well integrated into latent space. In order to assess the imputation performance of both models, we measured the relative errors of prediction (unseen modality) with respect to reconstruction (seen modality in the paired test set) (Methods section 4.4.6). This relative performance was similar for both multiDGD and MultiVI, although multiDGD shows a greater variance (Supplementary Table 4). However, the absolute prediction and reconstruction performances of multiDGD for ATAC data are still superior to those of MultiVI (Supplementary Table 5).

#### 2.5.2 Integrating unseen batches without architectural surgery

A novel feature of the DGD is its capability to find representations for previously unseen data. This can simply be unobserved cells from the seen covariates, but also completely new data from unobserved covariates. The latter is possible thanks to the probabilistic modelling of both the desired ‘molecular’ and covariate components of the representation. We explore the quality of representations and predictions for unseen data by applying the leave-one-out method to train the model. For each batch in the human bone marrow data (defined as the site the data was processed at), we train a multiDGD instance on the training samples of all other batches, providing us with four models. We evaluate these models on their test performances in terms of prediction errors relative to the model trained on all batches. In Fig. 4A, we see a marginal increase in the prediction loss of unseen batches as expected, but overall prediction performance is on par with the model trained on all batches (Fig. 4B) and the unseen batch samples are well integrated into the latent space (Fig. 4D). So far, unseen covariates have been integrated with approaches such as architectural surgery (scArches [28]). We include a comparison to scArches applied to MultiVI in the same scheme. However, due to the need for a fine tuning set, we run scArches on the training portion of the held-out batch, in order to keep the test set independent. This, of course, gives MultiVI+scArches an advantange of additional data. For MultiVI+scArches, overall reconstruction error decreases compared to the MultiVI model trained on all batches, highlighting the nature of fine-tuning in scArches. Absolute performance metrics, however, are still inferior to multiDGD (Fig. 4C) and integration into latent space is equivalent (Figure 4D and Supplementary Figure 14), making post-hoc fine tuning obsolete.

**Fig. 4.**
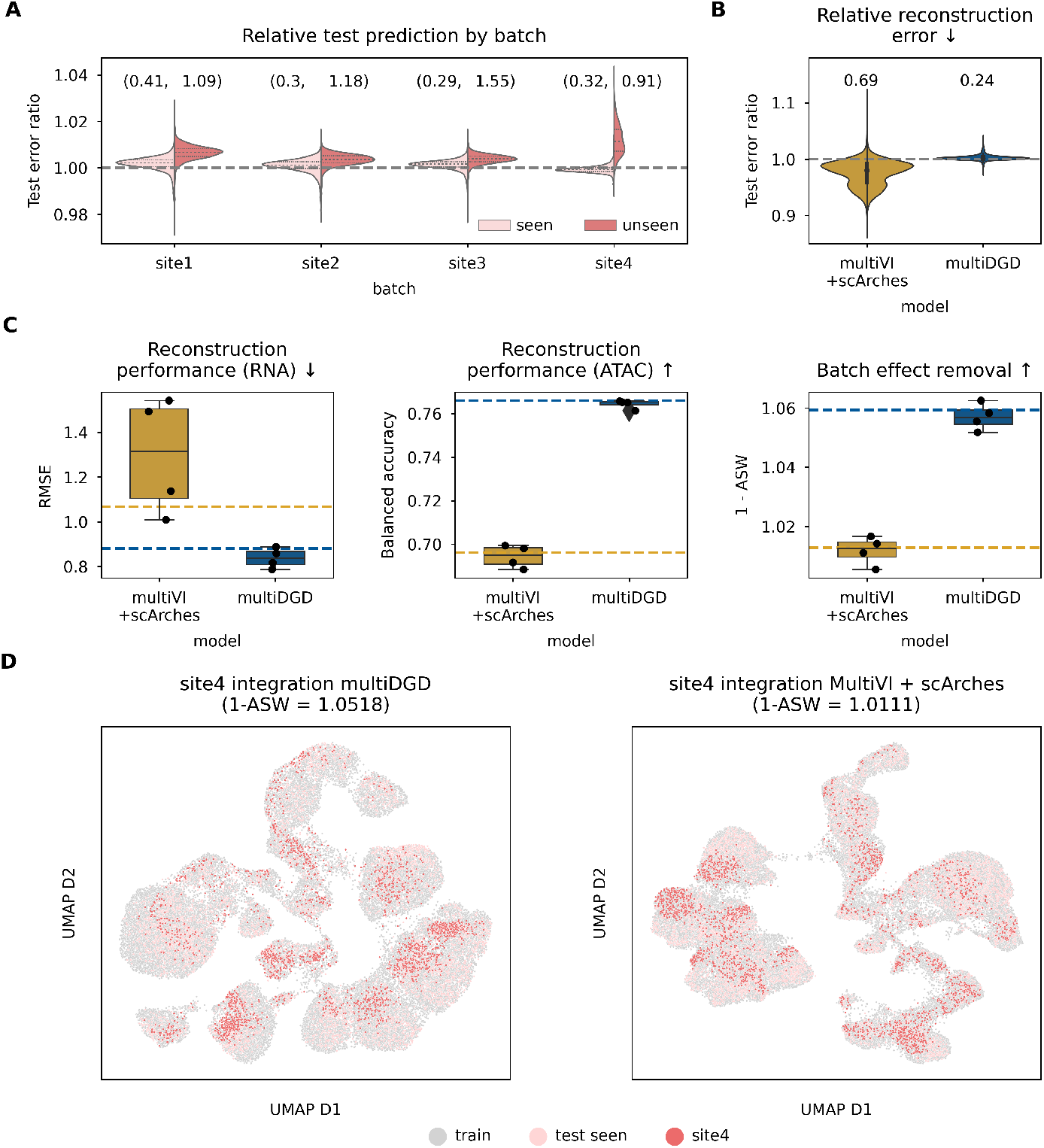
Model fine-tuning is no longer needed to predict unseen covariates. All related experiments were performed on the bone marrow data set. For each site (presenting the batches), one model was trained that excluded the site from training. This resulted in four models. Comparisons are done on test predictions with respect to the model trained on all sites (‘full’). scArches was applied with the left-out site from the train set to leave the test set independent. This results in a fine-tuned MultiVI model using all training data. **A)** Split violin plot of relative test errors for each multiDGD model wrt. the ‘full’ model. The relative errors are colored by whether the site had been included in training (seen) or not (unseen). **B)** Comparison of relative test errors for multiDGD and MultiVI fine-tuned with scArches. **A-B)** text above violins and boxes present the Kullback-Leibler divergences between the original test errors and the ones derived from the models trained on the incomplete data. **C)** Absolute performance comparison of multiDGD and MultiVI. From left to right: Reconstruction performances of multiDGD and MultiVI+scArches for RNA (I) and ATAC (II) data. Batch effect removal (III). Dashed lines present the original model performances for training on the full set. Box plots present the range of the performances of the four models from the leave-one-out scheme, which are shown as black dots. **D)** UMAP visualizations of latent spaces for multiDGD and MultiVI colored by the sample type. Samples were either taken from the train set or from the test set, which is distinguished by whether the batch was seen during training (seen) or not (unseen).

### 2.6 Gene-to-peak association with *in silico* perturbation

An emergent property of multiDGD is the learned connectivity between gene expression and chromatin accessibility data. We can use this to perform *in silico* predictions of where chromatin accessibility is associated with the expression of a given gene or set of genes. As depicted in Fig. 5A, we can compute gradients in latent space in the direction of perturbing a given gene 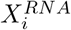 in data space. For every cell used, we have the original representation and a representation after one step of perturbation (*Z*^*KO*^). From these representations, we predict the perturbed sample and calculate the differences in prediction 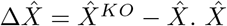 refers to model predictions.

**Fig. 5.**
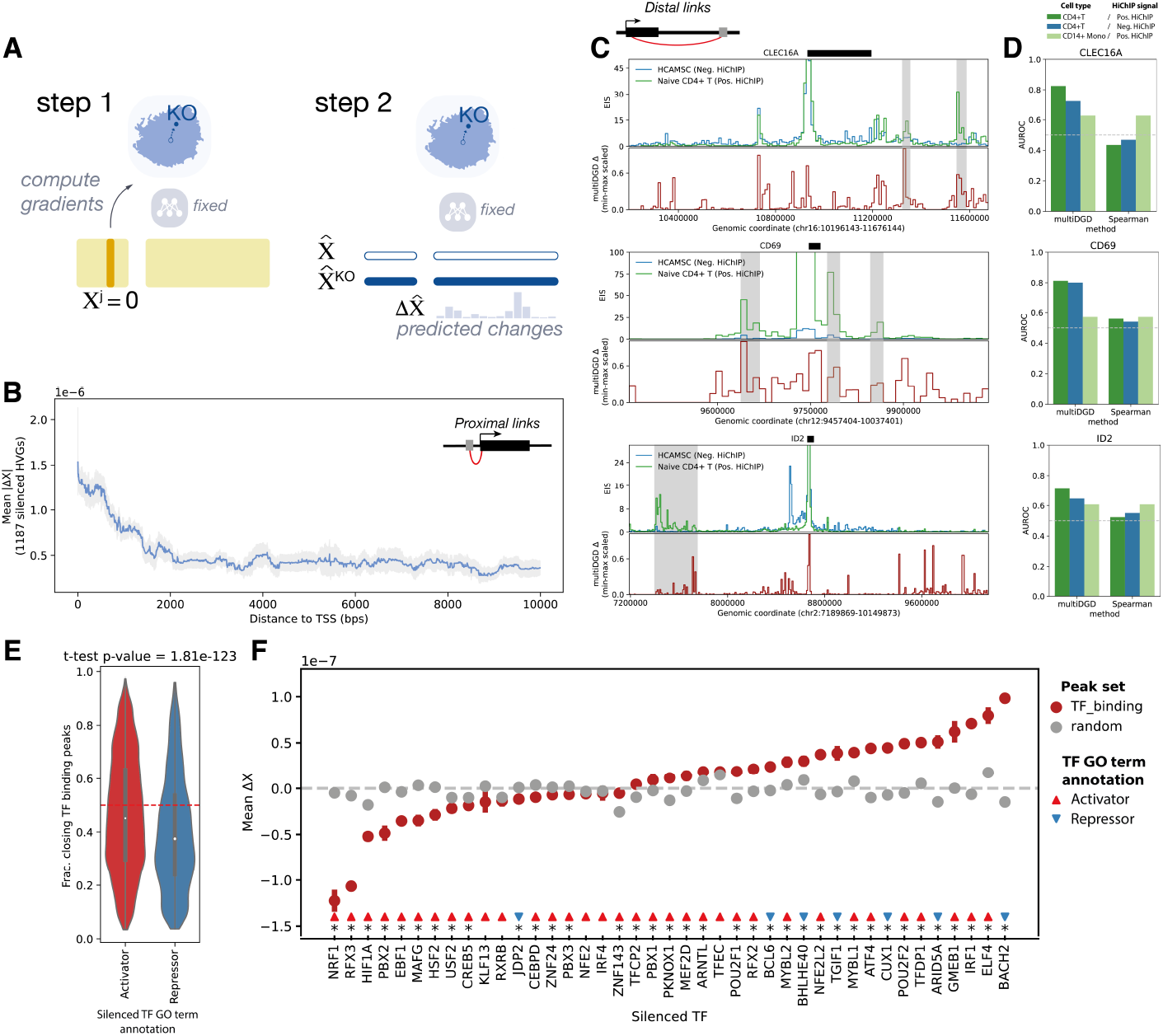
Prediction of association between gene expression and peak accessibility.. **A)** Schematic of gene-peak association prediction with *in silico* perturbation. **B)** Mean absolute effect of perturbation on peak accessibility (Δ*X*, y-axis) within 10 kb window around transcription start site (TSS) of silenced gene. The rolling average and 95% confidence interval of perturbation effect over the distance to TSS is shown for silencing of 862 highly variable genes (HVGs), with window size of 100 bps. **C)** Comparison of HiChIP signal around 3 genes (CLEC16A, CD69, ID2) with Δ*X* from silencing of the gene in naive CD4+T cells. For each gene, the top track shows the Enhancer Interaction Score (EIS) calculated from HiChIP data using the gene promoter as viewpoint (see methods), for HiChIP on primary naive CD4+ T cells (Positive HiChIP, green) and on the muscle cell line HCAMSC (Negative control HiChIP, blue). The bottom track (red line) shows the scaled Δ*X* for the prediction of peaks associated with expression of the gene of interest. The location of the transcript for the gene of interest is shown on top of each plot. T-cell-specific enhancer regions are highlighted in grey. **D)** Barplots of Area under the Receiver Operating Characteristic Curve (AUROC, y-axis) for prediction of HiChIP enhancer regions from Δ*X* or Spearmann correlation between gene expression and peak accessibility in the selected cell type (x-axis). We show AUROC for predicted associations in bone marrow CD4+ T cells of CD4+ T cell HiChIP signal (dark green) or HCAMSC HiChIP signal (blue), and for predicted associations in CD14+ monocytes of CD4+ T cell HiChIP signal (light green). **E)** Violin plots of fractions of closing peaks (Δ*X <* 0) for *in silico* silencing of transcription factors (TFs) annotated as activators (red) or repressors (blue). The black box and white point denote respectively the interquartile range and median of the distribution. The p-value for a 2-sample t-test comparing the mean of the distributions is shown. **F)** Mean Δ*X* in response to TF silencing for 34 predicted activators and 7 annotated repressors. We show the mean effect for all affected cells over a set of 10,000 peaks containing TF binding motifs (red points) or over a set of 10,000 peaks sampled at random amongst the peaks containing at least one TF binding motif (grey points) with the standard error of the mean. Asterisks denote TF perturbations for which the difference in mean effect between TF binding peaks and random peaks is significant (2-sample T-test p-value *<* 0.01).

First, we evaluated the ability of this *in silico* perturbation method to recover the association between gene expression and accessibility around the transcription start sites (TSS). When silencing a set of highly variable genes in the bone marrow data set, we observe significantly higher mean 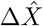 at peaks in the proximity of the TSS as expected. 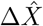 then gradually flattens for peaks over 2000 base pairs away (Figure 5B).

Next, we investigated whether we could recover associations between genes and peaks overlapping distal enhancers. As ground-truth data for gene-enhancer interaction, we used H3K27ac HiChIP data measuring physical contacts between active chromatin and promoters in primary CD4+T cells [39]. We predicted the effect on chromatin openness from silencing three genes with CRISPR-activation-validated T-cell-specific enhancers captured by HiChIP (CLEC16A, CD69, ID2) [39]. We observed high perturbation effect in naive CD4+T cells in several regions with HiChIP evidence (Figure 5C), which were not captured by HiChIP in a muscle cell line (negative control for cell-type-specific interactions). In addition, the enhancer prediction accuracy was significantly lower when considering perturbation effect in a different cell type (CD14+ monocytes) (Figure 5D). These results suggest that multiDGD is capturing cell-type-specific enhancer-gene links. In all cases, multiDGD prediction outperformed gene-peak associations derived from Spearmann correlation of gene expression and peak accessibility, where performance was close to random (Figure 5D, Supplementary Figure 15). We also observed instances of high 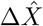 in absence of HiChIP signal (Figure 5). While these might be false positives driven by noise in the scATAC data, it is possible that multiDGD could be capturing indirect effects of gene expression on accessibility.

Finally, we sought to leverage *in silico* predictions with multiDGD to investigate the correspondence between the expression of transcription factors (TFs) and their effect on accessibility of peaks containing their DNA binding motifs. We tested the effect of silencing 41 TFs on chromatin accessibility, using a categorization of “activators” or “repressors” from Gene Ontology (GO) terms [40]. When measuring the perturbation effects at peaks containing TF binding motifs, we found that silencing activator TFs tends to lead to significantly higher fractions of closing peaks compared to silencing of annotated repressors (Figure 5E). For 36 out of 41 TF perturbations, we found a significant difference in mean 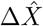 between peaks containing TF binding motifs and matched random peaks (T-test p-value *<* 0.01). However, we observe broad variation in perturbation effects for different TFs. We measure the strongest chromatin closing effects in response to silencing (indicating a positive correlation between accessibility and expression) for TFs with reported activation effects on a broad set of metabolic genes involved in stress response, including NRF1 [41] and HIF1A [42]. For about one third of the TFs annotated as activators based on GO terms, we detected perturbation effects at TF binding peaks consistent with repressive activity (i.e. chromatin opening upon TF silencing). In several cases these discrepancies could be explained by conflicting GO terms, where the same TF is reported to have both activator and repressor function in different cellular contexts (e.g. ELF4 [43], IRF1 [44], POU2F1-2 [45]). We frequently observe perturbation changes in both directions for this class of TFs, with mean 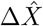 varying between cell types (Suppl. Figure 16B). PBX1, for example, is generally regarded as a transcriptional activator in the context of cancer, but can play a dual role in hematopoiesis [46]. In our data this factor is specifically expressed in HSCs and erythroid progenitors (Suppl. Figure 16A). While its *in silico* silencing leads to chromatin closing in most cell types, we predict chromatin activation in HSCs and granulocytic-myeloid progenitors (G/M prog, GMP). This is in line with evidence from mouse studies suggesting that PBX1 KO leads to premature derepression of GMP transcripts in myeloid progenitors [46]. These results further support the idea that multiDGD is learning cell-type-specific patterns of regulation and that coupling the generative model with *in silico* perturbation can be used to interpret the interplay between molecular features in cells.

## 3 Discussion

We present multiDGD, a novel generative model for single-cell multi-omics data. We demonstrate its use as a tool for dimensionality reduction and cross-modality prediction, but also new functionalities. Firstly, it enables the integrated modeling of unseen batches without the need for fine tuning methods such as scArches [28]. Secondly, it contains a built-in analysis of gene-peak associations. This is possible due to the emergent properties of generative models, which enable us to combine integration and analysis of data in a single framework. We have thus focused on comparing to existing generative models for multi-omics data [8, 11, 20] in this work. Among these, however, only MultiVI provides functionalities such as batch effect removal, which is critical for our analysis. We show that multiDGD presents a strong improvement compared to MultiVI [8], which is based on VAEs and presents a popular generative model architecture used in single-cell analysis. Our model outperforms MultiVI in terms of data reconstruction, cross-modality prediction and cell type clustering. Part of the performance increase on modelling ATAC data may be due to the use of raw counts rather than binarised data, preserving more information [47]. However, we attribute much of the general performance increase in comparison to MultiVI to the more complex latent distribution and the removal of the encoder. The performance increase in modelling RNA and ATAC counts can also be seen in the prediction of unseen cells and of missing modalities, although the variance in prediction performance is higher for multiDGD. One potential reason for this could be the over-denoising in MultiVI, which would also contribute to generally lower performance.

Even though multiDGD also outperforms the Leiden algorithm on MultiVI latent dimensions in terms of cell type clustering and learns meaningful embeddings, sampling from the prior over component means in the initialization leads to a high variance in clustering performance. We are eager to investigate how to stabilise this behaviour. Potential directions are to initialise the component means from origin as well or to re-sample representations during training to avoid local minima.

As single-cell multi-omics profiling becomes more robust and accessible, models for analysis will need to efficiently handle multi-sample data sets. They will further have to be able to disentangle technical batch effects from biological differences between cell types. We have introduced probabilistic modelling of covariates for the DGD. Probabilistic modelling of covariates improves basal representations, successfully captures interpretable sample-specific differences, and enables the integration of unseen data from different categories without architectural surgery. We have demonstrated this application and show that multiDGD can easily predict an unseen covariate with nearly the same performance as if it had been trained on it, but without any fine tuning. We believe this feature will facilitate the construction and re-use of large multi-omics atlases.

Multi-omics data sets, however, are often still small compared to scRNA-seq data sets [1] . The number of genomic peaks frequently outnumbers the amount of sequenced cells. Here we show that multiDGD shows clear advantages in modelling small data sets with high-dimensional spaces compared to data-hungry VAEs. We envision these capabilities will be of great value in allowing us to consider genome-wide epigenetic profiles for targeted analyses of data subsets of interest, such as specific lineages.

The goals of multi-omics analysis of course go way beyond efficient and high-quality embedding of cells. What is really desired is to further our understanding of gene regulation. Since we can incorporate larger decoders in multiDGD compared to VAE-based methods, explaining non-linear relationships may become easier. The resulting reconstruction performance increase certainly enables more reliable analysis at feature level. We made use of this in the prediction of gene-peak associations based on *in silico* perturbations.

We demonstrated meaningful associations in both proximal and distal interactions, and showed that the model can capture the effect of activating and repressing transcription factors at DNA binding sites. We recognize that at the current state, this gene-peak linkage prediction is a proof of concept, with several open questions to be investigated. For example, the extent by which multiDGD captures direct or secondary interactions remains to be determined, and whether different types of association can be distinguished. Further investigation is needed to determine whether the magnitude of perturbation changes is meaningful, or whether it is the most sensible to model counts as fractions of the count depth, as it is commonly done, for this application. Nevertheless, our results emphasize the potential of generative models as tools to capture interactions between molecular layers.

Altogether, multiDGD provides a strong performance increase on data reconstruction and clustering, incorporates modelling of covariates and provides a unified framework for integration and analysis of genomic features. We see these features as a significant next step in the evolution of single-cell multi-omics modelling.

## 4 Materials and Methods

### 4.1 Data

This work makes use of single-cell multiome data sets from human bone marrow, human brain tissue and mouse gastrulation stages.

#### 4.1.1 Acquisition

The human bone marrow multiome data set from [31] was downloaded from NCBI GEO [48] under accession GSE194122 on September 12 2022. For human brain data, we used the annotated data from [32] The annotated mouse gastrulation set used was taken from [33].

#### 4.1.2 Preprocessing

Human bone marrow data was used directly without any preprocessing. It comprises counts for 13431 gene transcripts and 116490 chromatin accessibility peaks. The 69249 cells represent 22 different cell types and were sequenced at four different sites. The sites are here interpreted as different batches.

The raw human brain counts were collected into an AnnData object according to [10X HDF5 Feature Barcode Matrix Format](https://support.10xgenomics.com/single-cell-multiome-atac-gex/software/pipelines/latest/advanced/h5_matrices). This data set contains 3534 cells with a total of 274892 features. We performed feature selection by excluding features that were not present (meaning counts of zero) in at least one percent of all cells. The result were 15172 transcripts and 95677 peaks. For cell type annotation, we chose the ATAC cell type annotation with 16 different types. From this data, we used no batch annotation.

For the mouse gastrulation data, we again performed feature selection based on the percentage of cells. The original number of features were 32285 for transcripts and 192251 for peaks. We excluded features that were only present in five percent of the cells and arrived at 11792 gene expression features and 69862 chromatin accessibility features. The total number of cells in this data is 56861 with 37 different cell types. This data contains a temporal component and thus makes the definition of batches more difficult. Nevertheless, we chose the gastrulation stage as the batch and expect this variable to only be partially removed from the latent representation as cell type appearance is not independent of the stage.

#### 4.1.3 Data splits

In order to adequately compare model performances across methods and random seeds, we created data splits for training, validation and testing. All three data sets were randomly split into a train set comprising 80 % of the samples and validation and held-out test sets with 10 % of the samples each.

### 4.2 The model

multiDGD is an extension of the Deep Generative Decoder (DGD) [18] for single-cell multiomics data. The core model consists of a decoder and a parameterized distribution over latent space. This is presented by a Gaussian Mixture Model (GMM) here. Since there is no encoder, inference of latent representations is achieved by learning representations as trainable parameters. This process is described in detail in [18]. multiDGD additionally offers the option of disentangled covariate representations. For this purpose, multiDGD learns not only a set of representations *Z* and distribution parameters *ϕ*, but also 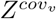 and 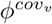 for every *v*th covariate. The corresponding graphical model is depicted in Supplementary Fig. 6. The following sections will describe the model and associated processes in more detail.

#### 4.2.1 Relevant notation

##### 4.2.2 Probabilistic formulation

The training objective is given by the joint probability *p*(*X, Z, θ, ϕ*) [18], which is maximized using Maximum a Posteriori estimation [18]. This can be decomposed into:

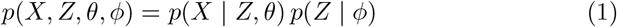

*p*(*X* | *Z, θ*) in this model is presented as the Negative Binomial distribution’s mass of the observed count *x*_*i*_ given the predicted mean count and a learned dispersion parameter *r*_*j*_ for each feature *j*

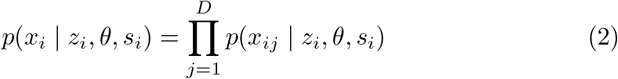

and

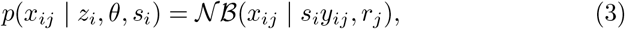

where *𝒩 ℬ* (*x*| *y, r*) is the negative binomial distribution. These equations are valid for each modality (RNA and ATAC) separately, as we have a total count *s* per modality.

The joint probability *p*(*X, Z, θ, ϕ*) that is maximized in the DGD [18] further contains the objective for the representation to follow the latent distribution, *p*(*Z* |*ϕ*). Since *ϕ* is a GMM, this results in the weighted multivariate Gaussian probability density

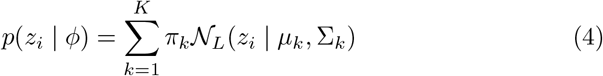

with K as the number of GMM components and *𝒩* _*L*_(*z*_*i*_ |*µ*, Σ) is a multivariate Gaussian distribution with dimension *L* (the latent dimension), mean vector *µ* and covariance matrix Σ.

For new data points the representation is found by maximizing *p*(*x*_*i*_ | *z*_*i*_, *θ, s*)*p*(*z*_*i*_|*ϕ*) only with respect to *z*_*i*_, as all other model parameters are fixed. More of this in section 4.2.8.

#### 4.2.3 Architecture

##### Decoder

The decoder in multiDGD is of hierarchical nature and will here be described in two sections: the shared network *θ*^*h*^ from latent space *Z* to the hidden state *H*, and the modality-specific networks *θ*^*mod*^ from *H* to their respective data spaces *X*^*mod*^. All layers in the networks consist of a linear layer followed by Rectified Linear Unit (ReLU) activation, except for the last layer in *θ*^*mod*^. The widths and depths of the networks are defined by hyperparameters described in Section 4.2.5. The activation of the last layer in *θ*^*mod*^ depends on the type of count scaling applied. Per default, the predicted normalized count means *y*^*mod*^ are scaled with the count depth *s*^*mod*^. The count depth presents the sum of all counts per modality. In this case, the predicted count means are achieved through softmax [49] activation of *y*^*mod*^. The probabilistic modelling of the counts and the corresponding objective function are described in the following section.

##### Count modelling

In multiDGD, counts of both gene expression and chromatin accessibility are modeled with Negative Binomial distributions (see Equation 3). For probabilistic modelling of outputs, we include ‘output modules’ which entail additional learned parameters and loss functions matching the probability distribution used. For the Negative Binomial output module, the necessary additional parameters are the feature-specific dispersion factors. For each feature in a given modality, we learn a dispersion factor to describe the shape of this individual feature’s distribution. The loss function in this module is given by the negative log probability mass function of the Negative binomial given an observed count. This provides us with the reconstruction loss of the given modality.

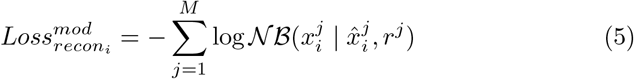

##### GMM (basal)

The Gaussian Mixture Model (GMM) presents the complex distribution over latent space in this model. It is a parameterized distribution which determines the shape of the latent space and is optimized in parallel to decoder and representation during training. The GMM consists of a set of *K* multivariate Gaussians with the same dimensionality as the corresponding representation. For the purpose of simplicity, we let multiDGD choose *K* based on the number of unique annotated cell types. This parameter is of course flexible and allows for tailored latent spaces depending on the desired clustering resolution. The objective for the representations is given as

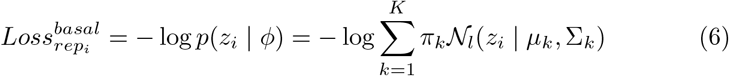

Trainable parameters include the means *µ* and covariances Σ of the components and the mixture coefficients *w*, which are transformed into mixture weights *π* through the softmax function. These parameters are in turn also learned with respective priors. The composition of the prior loss is given as follows.

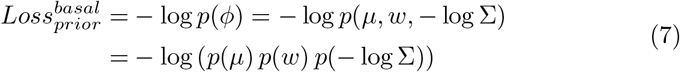

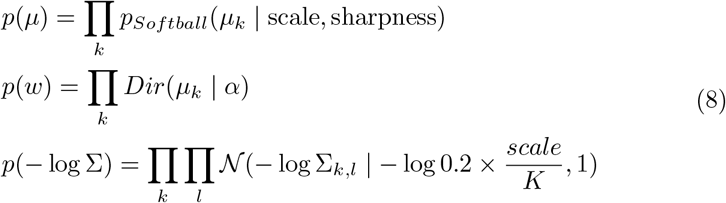

Altogether, these losses form the objective for both the representations and the GMM and will be referenced as the latent loss

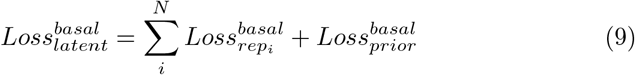

##### GMM (covariate)

The difference between the GMM for the basal latent space and the covariate space is merely the training scheme. As mentioned above, training for covariate representation and GMM is supervised. This results in a change in the objective as only probability densities of components assigned to a sample’s label are taken into account.

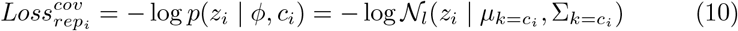

This means that the conditional probability *p*(*z*_*i*_ | *ϕ*) is solely dependent on the component with index identical to the numerical label *c*_*i*_ *∈*0, …, *C* with *C* as the number of unique covariate labels.

#### 4.2.4 Representations

Representations are treated as trainable parameters. However, they formally do not belong to the model architecture since they represent the low-dimensional embedding of data.

##### Representation (basal)

For each sample *x*_*i*_ with *i∈ N*, there exists one representation *z*_*i*_. The basal representations *Z*^*basal*^ represent the main embedding of data *X*, which aims to model the desired biological attributes of the data in low-dimensional space. As this structuring is unknown, *Z*^*basal*^ is inferred in an unsupervised setting. As described in [18], the representations are updated once per epoch with the gradients derived from reconstruction and distribution losses. Default initialization is given in Section 4.2.5.

##### Representation (covariate)

Covariates represent experimental variables we wish not to influence *Z*^*basal*^. In order to separate these influences, we model these attributes in distinct two-dimensional spaces *Z*^*cov*^. Here, it is necessary to follow a supervised training approach for successful disentanglement. This process is described in the corresponding section for the covariate GMM.

#### 4.2.5 Initialization and default parameters

The decoder contains two layers in the shared network and two in the modality-specific ones. All layers except the last one in modality-specific networks have 100 hidden units (*layer_width*). The last layer in a modality-specific network receives *max*(*layer_width*, 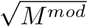) hidden units. Depth and width hyperparameters can of course be altered and should be considered depending on the number of samples and features available. Weights and biases are initialized per default using PyTorch’s Kaiming Uniform [50] method. In the Negative Binomial output module, dispersion parameters are initialized with a default value of 2.

Representations are generally initialized at origin, meaning they all start from zero vectors. One could also initialize from a pre-defined matrix, for example an *l*-dimensional Principal Component Analysis (PCA) or sampling from the prior. However, linear mappings are not always representative of the true underlying structure. In the default settings, the latent dimensionality *l* is set to 20, and covariate representations receive two dimensions.

The GMM is generally initialized with Softball prior scale 2 and hardness 5 and a Dirichlet *α* of 1. The prior over the covariance matrix Σ is defined by the number of mixture components as in [18] with 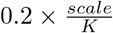. The GMM per default contains a single Gaussian. This setting is used if no observables are provided for the basal latent space. However, we do not recommend the single component as it will decrease the flexibility and complexity of the basal representation and will not provide the intrinsic clustering of the model. If an observable is given, the number of unique classes will be used as the number of components in the basal GMM *ϕ*. This observable is commonly the cell type label. For the covariate GMM *ϕ*^*cov*^, the number of components is equal to the number of unique categories in the covariate.

#### 4.2.6 Training

The general training algorithm remains as presented in [18], with an extension due to the covariate latent model and presence of multiple modalities.

##### Algorithm 1 Training

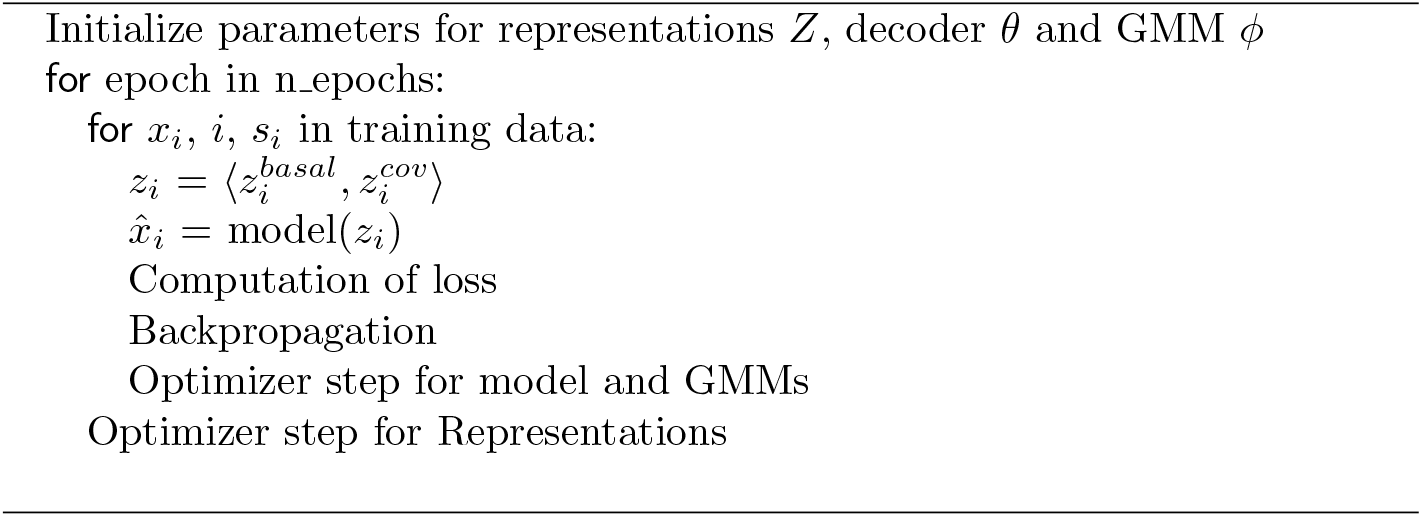

The training data is iterated over in mini batches with a default batch size of 128. Each set of parameters receives their on Adam [51] optimizer with betas (0.5, 0.7) and learning rates of 1*e−* 4 for the decoder and 1*e −*2 for representations and GMMs. As a proxy for the prior over *θ*, a weight decay of 1*e−* 4 is applied. The default maximum number of epochs is set to 1000, with early stopping applied at the earliest in epoch 50, taking into account the last 10 epochs.

The loss is presented as

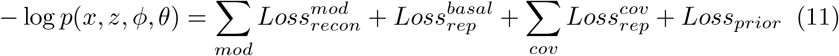

Positive definite parameters such as the Negative Binomial dispersion factors and the GMM covariances are learned as their logarithmic counterparts for numerical stability and enforcing the positive constraint.

#### 4.2.7 Validation

Validation is performed in parallel to training. Representations for the validation set are equally initialized at origin and optimized every epoch. In the validation loop, only the representation parameters of the validation set are updated, and covariate representations are inferred in an unsupervised manner.

#### 4.2.8 Testing and prediction of new data

With testing and predicting, we refer to the inference stages after the model parameters *θ* and *ϕ* have been trained and are regarded as frozen. The inference of representations for unseen data points is depicted in Fig. 1B. Firstly, the best mode (i.e. GMM component) is found for each sample. The best mode is given by the maximization of *p*(*x* | *z, θ, ϕ*) Π_*k*_ *p*(*z* | *ϕ*_*k*_) with respect to *k*. This step is the memory-critical process as for each new data point *X*_*m*_ with *m ∈M, K* losses have to be computed. In the case of present covariate models, this problem becomes combinatorial. In total, 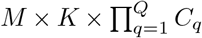 losses have to be computed, with *C*_*q*_ representing the number of covariate classes for covariate *q*.

After the best modes have been determined, the representations are optimized for a set number of steps, per default 10. This process is very fast and negligible compared to the total run time as long as the number of cells is in the thousands [18].

#### 4.2.9 Integrating unseen covariates

Because the covariate models are probabilistic, the method for integrating unseen covariate classes presented here works just like predicting new data. The unobserved covariate label might not have received its own distribution, but the model is capable to find the best covariate representation given its prior knowledge. One could see this as the unseen class being determined as a linear combination of the observed ones.

There is also the option of inferring an additional GMM component alongside the new covariate label. However, this is closer to fine-tuning as in scArches [28] and thus not discussed here, although we do plan to include this functionality in the future.

#### 4.2.10 Missing modality prediction

We again start from a trained model with all internal parameters (decoder and GMM) fixed. For the new samples, representations are initialized as described above. The only change to the simple data inference is that only the loss of the observed modality is used to infer representations. After inference, the predictions for all features are generated, so we get a completed picture of the sample.

#### 4.2.11 Internal clustering

The GMM is naturally equipped to cluster sample representations. Part of the objective calculation is to get a *K*-length vector for every representation containing the probability densities of said representation under each component *k ∈ K*. The argmax of this vector returns the index of the component with highest probability.

#### 4.2.12 Gene-to-peak association with *in silico* perturbation

Figure 1C depicts the mechanism of the gene2peak feature of multiDGD. Intuitively, we associate features to each other across modalities by predicting the effect of silencing a given gene or set of genes *X*^*j∈RNA*^ on the reconstructed peak accessibility profiles 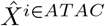. To simulate the effect of silencing gene *j* on chromatin accessibility, we consider cells in which *X*^*j∈RNA*^ *>* 0. Then for each cell *i* we generate a pseudo-expression profile *X*_*KO*_ where 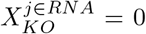 and compute the loss as usual. The loss is then backpropagated to get a gradient on *Z*^*basal*^. As a result, we have the original representation *Z*^*basal*^ and the perturbed representation 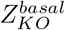. From both,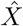 and 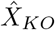 are predicted, and the perturbation changes 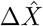 are computed as 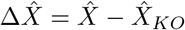.

### 4.3 Model search

#### 4.3.1 Hyperparameter optimization

Architectures and training parameters can vary strongly in both VAE and DGD. For a comparison of tools rather than Machine Learning methods, we chose the default settings for both MultiVI and multiDGD. The default settings for multiDGD are derived from design knowledge gained in [18]. Additional parameters such as model depth of the hierarchical DGD were found experimentally. Final default parameters are described in section 4.2.5 above. MultiVI models were trained with one, two and three layers in the encoder and decoder each. The best architecture was chosen for each individual data set. multiDGD does not have an encoder, but shared and modality-specific networks. We tested different depths of one to three layers for the shared and modality-specific feed-forward networks. A summary of our parameter search can be found in Supplementary Figure 7.

#### 4.3.2 Training

For each of the data sets included in this work, we train three instances of each method with random seeds 0, 37 and 8790.

#### 4.3.3 Model selection

For each data set, the best model from hyperparameter optimization was chosen based on the validation loss. This refers to the negative log density for multiDGD and the ELBO for MultiVI.

### 4.4 Performance evaluation

#### 4.4.1 Reconstruction

Reconstruction performance metrics were chosen based on their compatibility for both MultiVI and multiDGD. Expression count reconstructions could be compared directly as both methods model the counts with negative binomial distributions. We thus report RMSE and MAE of the test predictions.

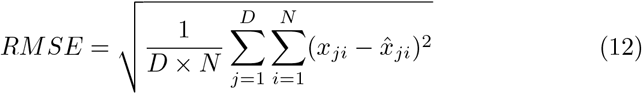

Chromatin accessibility is modelled differently in the two approaches. While multiDGD uses negative binomials, MultiVI models the peak counts as probabilities of being open. We thus binarized all predictions. As multiDGD does not use probabilities with a range from zero to one but models the actual counts, we calculated a prediction threshold of 0.2 for the binarization. After transforming the predictions, we compute the balanced accuracy as the average of sensitivity and specificity.

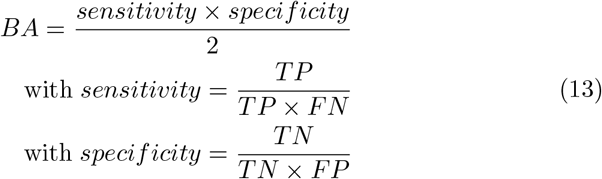

*TP, FN, TN, FP* present true positives, false negatives, true negatives and false positives, respectively. Positives are defined as ‘open’ regions (value = 1) and negatives as ‘closed’ (value = 0).

#### 4.4.2 Clustering

multiDGD can naturally cluster samples based on the probabilities of the GMM components for a given representation. This is not possible with MultiVI’s standard Gaussian prior, so it is common practice to perform Leiden [34] clustering on the latent space. We compare GMM and Leiden clustering through the Adjusted Rand Index (ARI) with respect to the cell type annotations. For a fair comparison, we adjust the Leiden algorithm such that it results in a similar number of clusters as the DGD, which is based on the number of unique cell types in the data. For bone marrow and brain data, the default scanpy Leiden parameters were used. For the gastrulation set, the resolution was set to 2. The ARI is the adjusted-for-chance version of the Rand index, which is related to clustering accuracy. This metric takes values between zero and one, with one representing perfect clustering according to the reference.

#### 4.4.3 Batch effect removal

The batch effect is measured as the Average Silhouette Width (ASW). It is given as

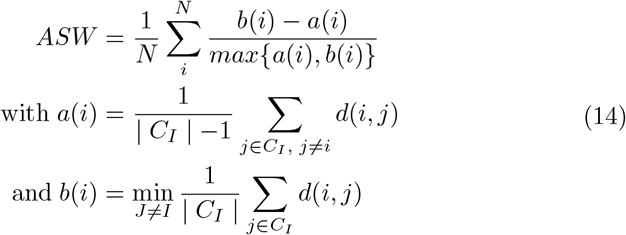

*a*(*i*) presents the mean distance between point *i* and all other points belonging to the same cluster. *d*(*i, j*) is the distance between points *i* and *j. b*(*i*) is the smallest mean distance of *i* to all points belonging to different clusters. This metric ranges between minus one and one. A value of one indicates a perfect clustering and a value of minus one indicates that samples would better fit into other clusters. For interpretability, we report 1 *− ASW* as the batch effect removal metric, where larger values indicate better performance.

#### 4.4.4 Data efficiency

In order to test model data efficiency, we created subsets of the largest data set in our study, with 1, 10, 25, 50 and 75 % of the training data. We trained both multiDGD and MultiVI instances on all these subsets with the hyperparameters determined for the full set and for the same three random seeds 0, 37 and 8790 as before. The performance of models on the subset is evaluated by the relative test losses, which we refer to as test loss ratios 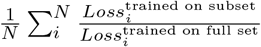 for every random seed.

#### 4.4.5 Feature efficiency

For this experiment we chose the mouse gastrulation data as it had previously been used with stringent feature selection (section 4.1.2) and offered the most additional features of all three data sets. The data with feature selection (features that were present in at least five percent of cells) contained 11792 genes and 69862 peaks. The full data set with all measured features is comprised of 32285 genes and 192251 peaks. We trained instances for both multiDGD and MultiVI on the full data set with random seed 0.

In order to assess in what way training on all features affected the models’ performances, we evaluated the reconstruction performances of RNA and ATAC data on the features previously selected for training (‘5%’).

#### 4.4.6 Modality prediction/imputation

The predictive performance is measured as the mean ratio of prediction error over reconstruction error 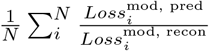. Prediction refers to the data generation of the missing modality in the unpaired samples. Data generation of the original, paired samples is described as reconstruction.

### 4.5 Gene-peak association

All gene-peak association predictions were performed on the test set of the bone marrow data (6925 cells).

For prediction of perturbation effects around transcription start sites (Figure 5B), we selected a sample of highly variable genes in the RNA data for the test set, using the method implemented in scanpy. We then ran the *in silico* silencing for all the sampled genes (as described in 4.2.12) and measured the mean perturbation effect on chromatin 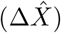 across all perturbed cells on peaks located within 10kb of the TSS of the silenced gene (using gene annotations from Ensembl v108). Of note, the mean perturbation effect across cells in TF binding sites is partially dependent on the total number of cells expressing the silenced gene in the test set (*R*^2^ = 0.45, p-value = 0.0029), suggesting that the estimates for perturbation effects might be more reliable with more support data.

For validation of gene-peak association predictions in distal enhancers (Figure 5C-D), we downloaded H3K27ac HiChIP data from primary T cells [39] from the Gene Expression Omnibus (GSE101498). Raw .hic files were converted to matrices of interaction signal between any two genomic bins of size 10kb using Juicebox tools [52], replicating the workflow and parameters described in [53]. We then computed the mean enhancer interaction signal (EIS) between 2 replicate samples for naive CD4+T cells and HCAMSC cells using as viewpoint the bins containing the promoter of the gene of interest. We calculated cell-type-specific gene-peak associations the absolute predicted change 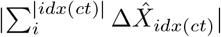 where *ct* stands for cell type and *idx*(*ct*) is the subset of the indices for this cell type. We evaluate the ability to recover enhancer-gene interactions from HiChIP data with ROC curve analysis, where we consider a genomic bin to be an enhancer region if the EIS is higher than the 75% quantile computed over the whole locus.

For the TF perturbation analysis (Figure 5E-F), we considered a list of transcription factors annotated as activators or repressors based on mining of GO terms by [40]. We identified peaks containing TF binding motifs using the JASPAR database (release 2022). We then restricted our analysis to TFs which had matches in less than 80% of all peaks and that were expressed in at least 250 cells in the test set. We computed 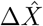 for silencing of each TF and computed the mean 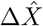 across 10k peaks sampled amongst the peaks containing TF binding motifs, and across 10k peaks sampled amongst all peaks containing at least one TF binding motif. This strategy to select the null ‘random’ set was used to exclude peaks with extremely sparse counts at distal intergenic locations, which might represent an unfair comparison for this analysis.

### 4.6 Visualization

We applied UMAP[54] to dimensionality reductions used in visualizations.

### 4.7 Software and Hardware

All code is written in Python (version *≥* 3.9.12) and executed on a cluster with x86 64 architecture and NVIDIA TITAN RTX and NVIDIA TITAN X GPUs. The machine learning framework used was PyTorch [55] version 1.10. Training progress was monitored and logged using weights and biases (wandb) [56]. MultiVI was used as part of the scvi-tools [57] package with version 0.19.0. For Cobolt and scMM, we applied version v1.0.1 and release 1, respectively. The scanpy [58] package was used for parts of the analysis.

## Acknowledgments

This publication is part of the Human Cell Atlas (www.humancellatlas.org/publications/). We acknowledge all the great discussions at the Sanger Institute regarding data generation, processing and analysis and the wonderful support from the Center for Health Data Science.

## Author contributions

A.K. contributed the theoretical foundation of the approach. A.K. contributed to the implementation of the approach. A.K. contributed to the writing of the manuscript.

E.D. contributed to the acquisition and preprocessing of data. E.D. contributed to the experimental designs in the work. E.D. contributed to the data analysis experiments. E.D. contributed to the writing of the manuscript. E.D. contributed to figures.

S.A.T. contributed to the experimental designs in the work. S.A.T. contributed to the writing of the manuscript.

V.S. contributed the theoretical foundation of the approach. V.S. contributed to the implementation of the approach. V.S. contributed to the acquisition and preprocessing of data. V.S. contributed to the experimental designs in the work. V.S. contributed to the writing of the manuscript. V.S. contributed the main experiments. V.S. contributed to figures.

## Code availability

The multiDGD code and package is made available here https://github.com/Center-for-Health-Data-Science/multiDGD.

The code to reproduce the presented results is available here https://github.com/Center-for-Health-Data-Science/multiDGD_paper.

## Data availability

In this work we made use of three publicly available data sets. These are a human bone marrow set [31] with accession number GSE194122, a mouse gastrulation set [33] with accession number GSE205117 and a human brain data set from [32] with accession number GSE162170. All processed data and trained model parameters have been deposited on Figshare (https://doi.org/10.6084/m9.figshare.23796198.v1).

## References

[1] Baysoy A, Bai Z, Satija R, Fan R. The technological landscape and applications of single-cell multi-omics;p. 1–19. Publisher: Nature Publishing Group. https://doi.org/10.1038/s41580-023-00615-w.

[2] Argelaguet R, Cuomo ASE, Stegle O, Marioni JC. Computational principles and challenges in single-cell data integration;p. 1– Bandiera abtest: a Cg type: Nature Research Journals Primary atype: Reviews Publisher: Nature Publishing Group Subject term: Computational biology and bioinformatics;Systems biology Subject term id: computational-biology-and-bioinformatics;systemsbiology. https://doi.org/10.1038/s41587-021-00895-7.

[3] Argelaguet R, Arnol D, Bredikhin D, Deloro Y, Velten B, Marioni JC, et al. MOFA+: a statistical framework for comprehensive integration of multi-modal single-cell data;21(1):111. https://doi.org/10.1186/s13059-020-02015-1.

[4] Stuart T, Butler A, Hoffman P, Hafemeister C, Papalexi E, Mauck WM, et al. Comprehensive Integration of Single-Cell Data;177(7):1888–1902.e21. Publisher: Elsevier. https://doi.org/10.1016/j.cell.2019.05.031.

[5] Welch JD, Kozareva V, Ferreira A, Vanderburg C, Martin C, Macosko EZ. Single-Cell Multi-omic Integration Compares and Contrasts Features of Brain Cell Identity;177(7):1873–1887.e17. https://doi.org/10.1016/j.cell.2019.05.006.

[6] Hao Y, Hao S, Andersen-Nissen E, Mauck WM, Zheng S, Butler A, et al. Integrated analysis of multimodal single-cell data;184(13):3573–3587.e29. Publisher: Elsevier. https://doi.org/10.1016/j.cell.2021.04.048.

[7] Singh R, Hie BL, Narayan A, Berger B. Schema: metric learning enables interpretable synthesis of heterogeneous single-cell modalities;22(1):131. https://doi.org/10.1186/s13059-021-02313-2.

[8] Ashuach T, Gabitto MI, Koodli RV, Saldi GA, Jordan MI, Yosef N. MultiVI: deep generative model for the integration of multimodal data. Nature Methods. 2023 Jun;p. 1–10. Publisher: Nature Publishing Group. https://doi.org/10.1038/s41592-023-01909-9.

[9] Hao Y, Stuart T, Kowalski MH, Choudhary S, Hoffman P, Hartman A, et al. Dictionary learning for integrative, multimodal and scalable single-cell analysis;p. 1–12. Publisher: Nature Publishing Group. phttps://doi.org/10.1038/s41587-023-01767-y.

[10] Ghazanfar S, Guibentif C, Marioni JC. Stabilized mosaic single-cell data integration using unshared features;p. 1–9. Publisher: Nature Publishing Group. https://doi.org/10.1038/s41587-023-01766-z.

[11] Gong B, Zhou Y, Purdom E. Cobolt: integrative analysis of multimodal single-cell sequencing data;22(1):351. https://doi.org/10.1186/s13059-021-02556-z.

[12] Luecken MD, Burkhardt DB, Cannoodt R, Lance C, Agrawal A, Aliee H, et al. A sandbox for prediction and integration of DNA, RNA, and protein data in single cells;.

[13] Eraslan G, Simon LM, Mircea M, Mueller NS, Theis FJ. Single-cell RNA-seq denoising using a deep count autoencoder;10(1):390. Number: 1 Publisher: Nature Publishing Group. https://doi.org/10.1038/s41467-018-07931-2.

[14] Lopez R, Regier J, Cole MB, Jordan MI, Yosef N. Deep generative modeling for single-cell transcriptomics;15(12):1053–1058. Number: 12 Primary atype: Research Publisher: Nature Publishing Group Subject term: Computational biology and bioinformatics;Computational models Subject term id: computationalbiology-and-bioinformatics;computational-models. https://doi.org/10.1038/s41592-018-0229-2.

[15] Xu C, Lopez R, Mehlman E, Regier J, Jordan MI, Yosef N. Probabilistic harmonization and annotation of single-cell transcriptomics data with deep generative models;17(1):e9620. Publisher: John Wiley & Sons, Ltd. https://doi.org/10.15252/msb.20209620.

[16] Lotfollahi M, Wolf FA, Theis FJ. scGen predicts single-cell perturbation responses;16(8):715. https://doi.org/10.1038/s41592-019-0494-8.

[17] Grønbech CH, Vording MF, Timshel PN, Sønderby CK, Pers TH, Winther O. scVAE: variational auto-encoders for single-cell gene expression data. Bioinformatics. 2020 Aug;36(16):4415–4422. https://doi.org/10.1093/bioinformatics/btaa293.

[18] Schuster V, Krogh A.: The Deep Generative Decoder: MAP estimation of representations improves modeling of single-cell RNA data.

[19] Lotfollahi M, Litinetskaya A, Theis FJ.: Multigrate: single-cell multiomic data integration.bioRxiv. Pages: 2022.03.16.484643 Section: New Results. Available from: https://www.biorxiv.org/content/10.1101/2022.03.16.484643v1.

[20] Minoura K, Abe K, Nam H, Nishikawa H, Shimamura T. A mixtureof-experts deep generative model for integrated analysis of single-cell multiomics data. Cell Reports Methods. 2021;1(5):100071. https://doi. org/https://doi.org/10.1016/j.crmeth.2021.100071.

[21] Cui H, Wang C, Maan H, Wang B.: scGPT: Towards Building a Foundation Model for Single-Cell Multi-omics Using Generative AI. bioRxiv. Pages: 2023.04.30.538439 Section: New Results. Available from: https://www.biorxiv.org/content/10.1101/2023.04.30.538439v1.

[22] Lopez R, Gayoso A, Yosef N. Enhancing scientific discoveries in molecular biology with deep generative models;16(9):e9198. Publisher: John Wiley & Sons, Ltd. https://doi.org/10.15252/msb.20199198.

[23] Kingma DP, Welling M.: Auto-Encoding Variational Bayes. arXiv. ArXiv:1312.6114 [cs, stat]. Available from: http://arxiv.org/abs/1312.6114.

[24] Luecken MD, Büttner M, Chaichoompu K, Danese A, Interlandi M, Mueller MF, et al. Benchmarking atlas-level data integration in single-cell genomics;p. 2020.05.22.111161. Publisher: Cold Spring Harbor Laboratory Section: New Results. https://doi.org/10.1101/2020.05.22.111161.

[25] Suo C, Dann E, Goh I, Jardine L, Kleshchevnikov V, Park JE, et al. Mapping the developing human immune system across organs;376(6597):eabo0510. Publisher: American Association for the Advancement of Science. https://doi.org/10.1126/science.abo0510.

[26] Eraslan G, Drokhlyansky E, Anand S, Fiskin E, Subramanian A, Slyper M, et al. Single-nucleus cross-tissue molecular reference maps toward understanding disease gene function;376(6594):eabl4290. Publisher: American Association for the Advancement of Science. https://doi.org/10.1126/science.abl4290.

[27] Sikkema L, Ramírez-Suástegui C, Strobl DC, Gillett TE, Zappia L, Madissoon E, et al. An integrated cell atlas of the lung in health and disease;29(6):1563–1577. Number: 6 Publisher: Nature Publishing Group. https://doi.org/10.1038/s41591-023-02327-2.

[28] Lotfollahi M, Naghipourfar M, Luecken MD, Khajavi M, Büttner M, Wagenstetter M, et al. Mapping single-cell data to reference atlases by transfer learning. Nature Biotechnology. 2022 Jan;40(1):121–130. Number: 1 Publisher: Nature Publishing Group. https://doi.org/10.1038/s41587-021-01001-7.

[29] Lance C, Luecken MD, Burkhardt DB, Cannoodt R, Rautenstrauch P, Laddach A, et al.: Multimodal single cell data integration challenge: results and lessons learned [preprint]. Available from: http://biorxiv.org/lookup/doi/10.1101/2022.04.11.487796.

[30] Schuster V, Krogh A. A Manifold Learning Perspective on Representation Learning: Learning Decoder and Representations without an Encoder. Entropy. 2021;23(11). https://doi.org/10.3390/e23111403.

[31] Luecken M, Burkhardt D, Cannoodt R, Lance C, Agrawal A, Aliee H, et al. A sandbox for prediction and integration of DNA, RNA, and proteins in single cells. In: Vanschoren J, Yeung S, editors. Proceedings of the Neural Information Processing Systems Track on Datasets and Benchmarks. vol. 1; 2021. Available from: https://datasets-benchmarks-proceedings.neurips.cc/paper/2021/file/158f3069a435b314a80bdcb024f8e422-Paper-round2.pdf.

[32] Trevino AE, Müller F, Andersen J, Sundaram L, Kathiria A, Shcherbina A, et al. Chromatin and gene-regulatory dynamics of the developing human cerebral cortex at single-cell resolution. Cell. 2021 Sep;184(19):5053–5069.e23. https://doi.org/10.1016/j.cell.2021.07.039.

[33] Argelaguet R, Lohoff T, Li JG, Nakhuda A, Drage D, Krueger F, et al.: Decoding gene regulation in the mouse embryo using single-cell multiomics. bioRxiv. Pages: 2022.06.15.496239 Section: New Results. Available from: https://www.biorxiv.org/content/10.1101/2022.06.15.496239v2.

[34] Traag VA, Waltman L, van Eck NJ. From Louvain to Leiden: guaranteeing well-connected communities. Scientific Reports. 2019 Mar;9(1):5233. Number: 1 Publisher: Nature Publishing Group. https://doi.org/10.1038/s41598-019-41695-z.

[35] Buettner F, Natarajan KN, Casale FP, Proserpio V, Scialdone A, Theis FJ, et al. Computational analysis of cell-to-cell heterogeneity in singlecell RNA-sequencing data reveals hidden subpopulations of cells. Nature Biotechnology. 2015 Feb;33(2):155–160. Number: 2 Publisher: Nature Publishing Group. https://doi.org/10.1038/nbt.3102.

[36] Bardot ES, Hadjantonakis AK. Mouse gastrulation: Coordination of tissue patterning, specification and diversification of cell fate. Mechanisms of Development. 2020;163:103617. https://doi.org/https://doi.org/10.1016/j.mod.2020.103617.

[37] Cremer C, Li X, Duvenaud D. Inference Suboptimality in Variational Autoencoders. arXiv:180103558 [cs, stat]. 2018 May;ArXiv: 1801.03558.

[38] Heumos L, Schaar AC, Lance C, Litinetskaya A, Drost F, Zappia L, et al. Best practices for single-cell analysis across modalities;p. 1–23. Publisher: Nature Publishing Group. https://doi.org/10.1038/s41576-023-00586-w.

[39] Mumbach MR, Satpathy AT, Boyle EA, Dai C, Gowen BG, Cho SW, et al. Enhancer connectome in primary human cells identifies target genes of disease-associated DNA elements;49(11):1602–1612. Number: 11 Publisher: Nature Publishing Group. https://doi.org/10.1038/ng.3963.

[40] Domcke S, Hill AJ, Daza RM, Cao J, O’Day DR, Pliner HA, et al. A human cell atlas of fetal chromatin accessibility;370(i6518). Publisher: American Association for the Advancement of Science Section: Research Article. https://doi.org/10.1126/science.aba7612.

[41] Ruvkun G, Lehrbach N. Regulation and Functions of the ER-Associated Nrf1 Transcription Factor;15(1):a041266. Company: Cold Spring Harbor Laboratory Press Distributor: Cold Spring Harbor Laboratory Press Institution: Cold Spring Harbor Laboratory Press Label: Cold Spring Harbor Laboratory Press Publisher: Cold Spring Harbor Lab. https://doi.org/10.1101/cshperspect.a041266.

[42] Corcoran SE, O’Neill LAJ. HIF1 and metabolic reprogramming in inflammation;126(10):3699–3707. Publisher: American Society for Clinical Investigation. https://doi.org/10.1172/JCI84431.

[43] Suico MA, Shuto T, Kai H. Roles and regulations of the ETS transcription factor ELF4/MEF;9(3):168–177. https://doi.org/10.1093/jmcb/mjw051.

[44] Fragale A, Gabriele L, Stellacci E, Borghi P, Perrotti E, Ilari R, et al. IFN regulatory factor-1 negatively regulates CD4+ CD25+ regulatory T cell differentiation by repressing Foxp3 expression;181(3):1673–1682. https://doi.org/10.4049/jimmunol.181.3.1673.

[45] Hwang SS, Kim LK, Lee GR, Flavell RA. Role of OCT-1 and partner proteins in T cell differentiation;1859(6):825–831. https://doi.org/10.1016/j.bbagrm.2016.04.006.

[46] Ficara F, Crisafulli L, Lin C, Iwasaki M, Smith KS, Zammataro L, et al. Pbx1 restrains myeloid maturation while preserving lymphoid potential in hematopoietic progenitors;126(14):3181–3191. https://doi.org/10.1242/jcs.125435.

[47] Martens LD, Fischer DS, Theis FJ, Gagneur J.: Modeling fragment counts improves single-cell ATAC-seq analysis. bioRxiv. Pages: 2022.05.04.490536 Section: New Results. Available from: https://www.biorxiv.org/content/10.1101/2022.05.04.490536v1.

[48] Edgar R, Domrachev M, Lash AE. Gene Expression Omnibus: NCBI gene expression and hybridization array data repository. Nucleic Acids Research. 2002 01;30(1):207–210. https://doi.org/10.1093/nar/30.1.207. https://academic.oup.com/nar/article-pdf/30/1/207/9901036/300207.pdf.

[49] Boltzmann L, Hasenöhrl F. Studien über das Gleichgewicht der lebendigen Kraft zwischen bewegten materiellen Punkten; 2012. .

[50] He K, Zhang X, Ren S, Sun J.: Delving Deep into Rectifiers: Surpassing Human-Level Performance on ImageNet Classification. arXiv. ArXiv:1502.01852 [cs] version: 1. Available from: http://arxiv.org/abs/1502.01852.

[51] Kingma DP, Ba J.: Adam: A Method for Stochastic Optimization. Cite arxiv:1412.6980. Published as a conference paper at the 3rd International Conference for Learning Representations, San Diego, 2015. Available from: http://arxiv.org/abs/1412.6980.

[52] Durand NC, Shamim MS, Machol I, Rao SSP, Huntley MH, Lander ES, et al. Juicer Provides a One-Click System for Analyzing Loop-Resolution Hi-C Experiments;3(1):95–98. Publisher:Elsevier. https://doi.org/10.1016/j.cels.2016.07.002.

[53] Granja JM, Klemm S, McGinnis LM, Kathiria AS, Mezger A, Corces MR, et al. Single-cell multiomic analysis identifies regulatory programs in mixed-phenotype acute leukemia;p. 1–8. https://doi.org/10.1038/s41587-019-0332-7.

[54] McInnes L, Healy J, Saul N, Grossberger L. UMAP: Uniform Manifold Approximation and Projection. The Journal of Open Source Software. 2018;3(29):861.

[55] Paszke A, Gross S, Massa F, Lerer A, Bradbury J, Chanan G, et al. PyTorch: An Imperative Style, High-Performance Deep Learning Library. In: Wallach H, Larochelle H, Beygelzimer A, d’Alché-Buc F, Fox E, Garnett R, editors. Advances in Neural Information Processing Systems 32. Curran Associates, Inc.; 2019. p. 8024–8035. Available from: http://papers.neurips.cc/paper/9015-pytorch-an-imperative-style-high-performance-deep-learning-library.pdf.

[56] Biewald L.: Experiment Tracking with Weights and Biases. Software available from wandb.com. Available from: https://www.wandb.com/.

[57] Gayoso A, Lopez R, Xing G, Boyeau P, Valiollah Pour Amiri V, Hong J, et al. A Python library for probabilistic analysis of single-cell omics data. Nature Biotechnology. 2022 Feb;40(2):163–166. Number: 2 Publisher: Nature Publishing Group. https://doi.org/10.1038/s41587-021-01206-w.

[58] Wolf FA, Angerer P, Theis FJ. SCANPY: large-scale single-cell gene expression data analysis. Genome Biology. 2018 Feb;19(1):15. https://doi.org/10.1186/s13059-017-1382-0.

