## Supplementary materials for "multiDGD: A versatile deep generative model for multi-omics data"

### Supplementaries

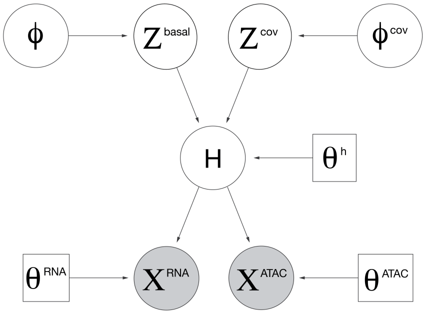


**Supplementary Figure 6 Graphical model of multiDGD.** Circles represent variables and boxes represent non-probabilistic parameters. Filled circles show observable variables. H represents the hidden intermediate state that is the output from $\theta^h$ and input to both $\theta^{RNA}$ and $\theta^{ATAC}$.

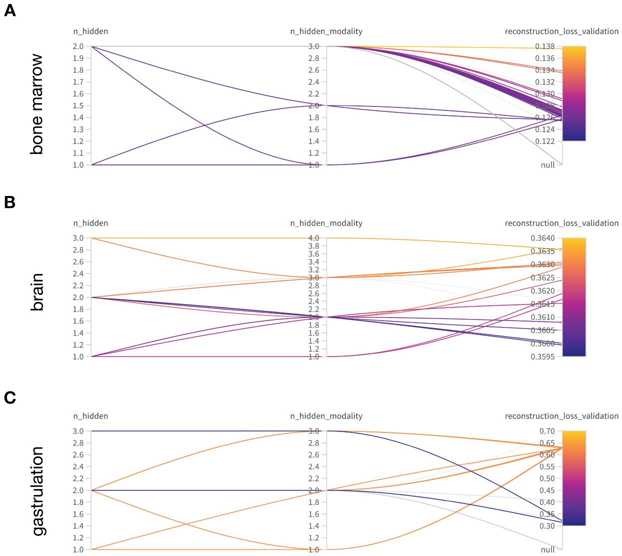


**Supplementary Figure 7 Hyperparameter optimization results of multiDGD decoder depths in weights and biases.** Each panel shows a parallel coordinate plot colored by the validation reconstruction loss. Hyperparameters of interest were the number of layers for the shared and modality-specific decoders. A-C shows results for the human bone marrow, human brain and mouse gastrulation data set, respectively.

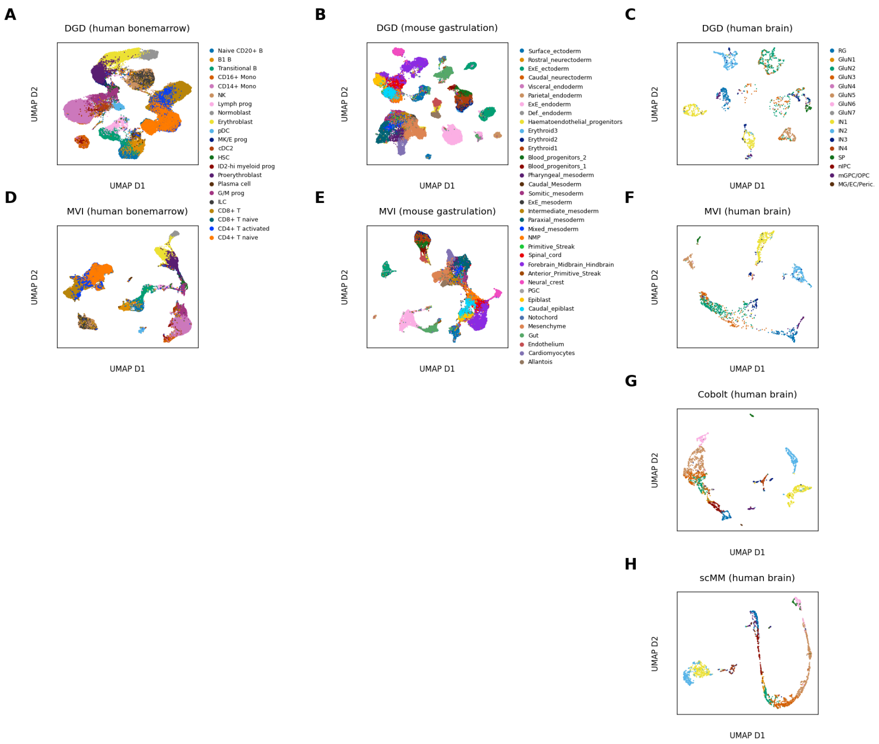


**Supplementary Figure 8 UMAP visualizations of model embeddings**. Titles indicate model and data set. **A-C)** Basal latent embeddings derived from multiDGD. UMAPs were created with default number of neighbors 15 and a minimum distance of 0.5. **D-F)** Basal latent embeddings derived from MultiVI. UMAPs were created with default number of neighbors 15 and a default minimum distance of 0.1. **G-H)** Latent embeddings for the human brain set from Cobolt and scMM, respectively. UMAPs were created with the same parameters as for MultiVI.

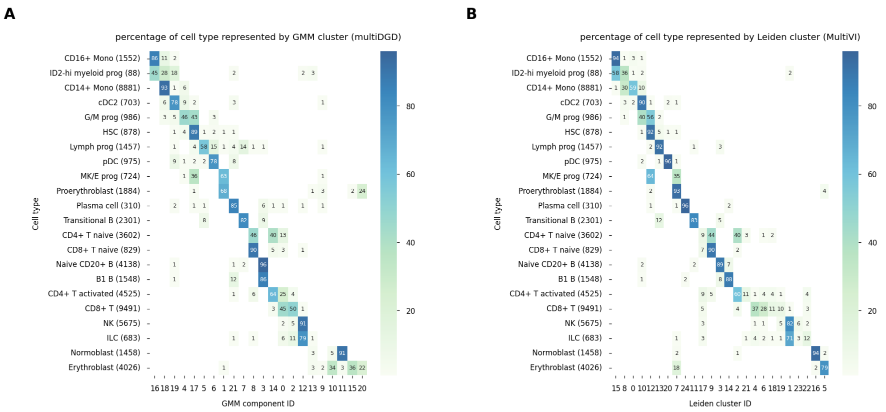


**Supplementary Figure 9 Clustering as percentage of cell types in the human bone marrow set as in 2F**. The x axis presents the component IDs, the y axis the annotated cell types. Numbers inside the heatmap indicate the percentage of samples in a cell type assigned to a given component. **A)** multiDGD. **B)** MultiVI.

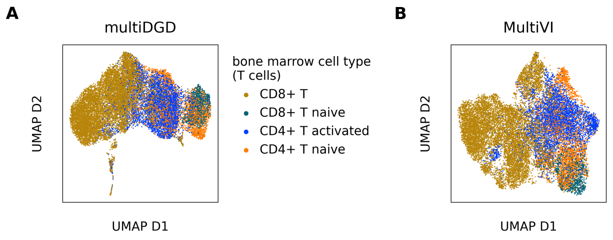


**Supplementary Figure 10 UMAP visualizations of T cells from the human bone marrow data set for multiDGD and MultiVI**. The depicted cells are from the training set. **A)** multiDGD. **B)** MultiVI.

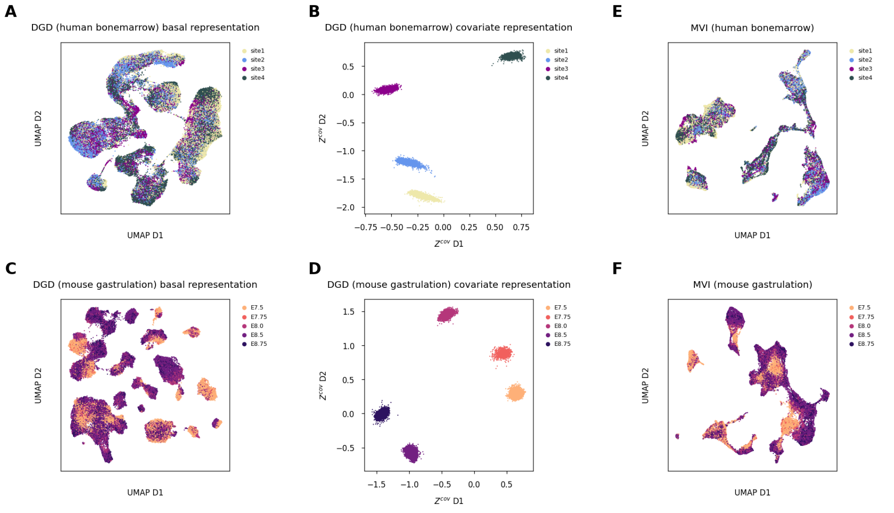

**Supplementary Figure 11 Visualizations of model embeddings colored by covariates.** Titles indicate model, data and type of representation depicted. **A)** Basal latent representation of the human bone marrow data set derived from multiDGD visualized as the first two dimensions of a UMAP. **B)** Two-dimensional covariate representation of the human bone marrow data set. **C-D)** Same as A and B for the mouse gastrulation data, respectively. **E-F)** UMAP visualizations of the MultiVI basal representations for the human bone marrow and mouse gastrulation data, respectively.

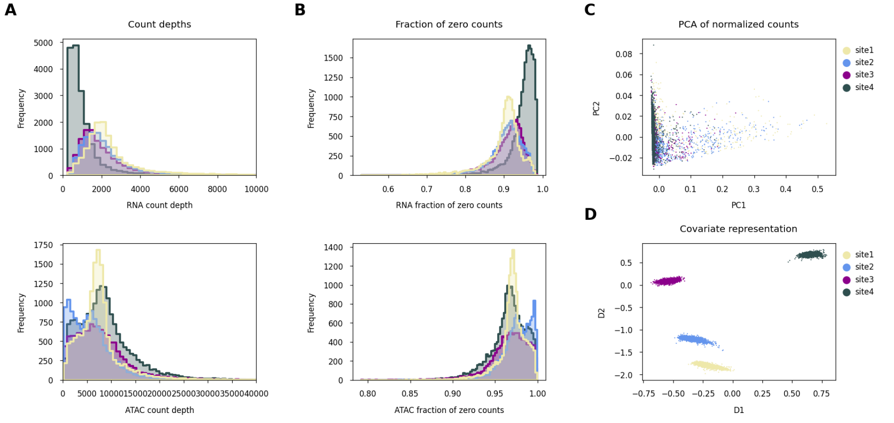

**Supplementary Figure 12 Distributional differences between sequencing sites in the human bone marrow data (train set)**. All subplots are colored by the site covariate (in the text referred to as the 'batch'). **A)** Histogram of count depths of RNA (top) and ATAC (bottom) cells. **B)** Histogram of fraction of zero counts of RNA (top) and ATAC (bottom) cells.**C)** Scatter plot of the first two principal components of the combined normalized modalities. Normalization was achieved by dividing RNA and ATAC counts by their respective count depths (after adding 1 to avoid collapse) and subsequent log-scaling. **D)** Batch representation of multiDGD.

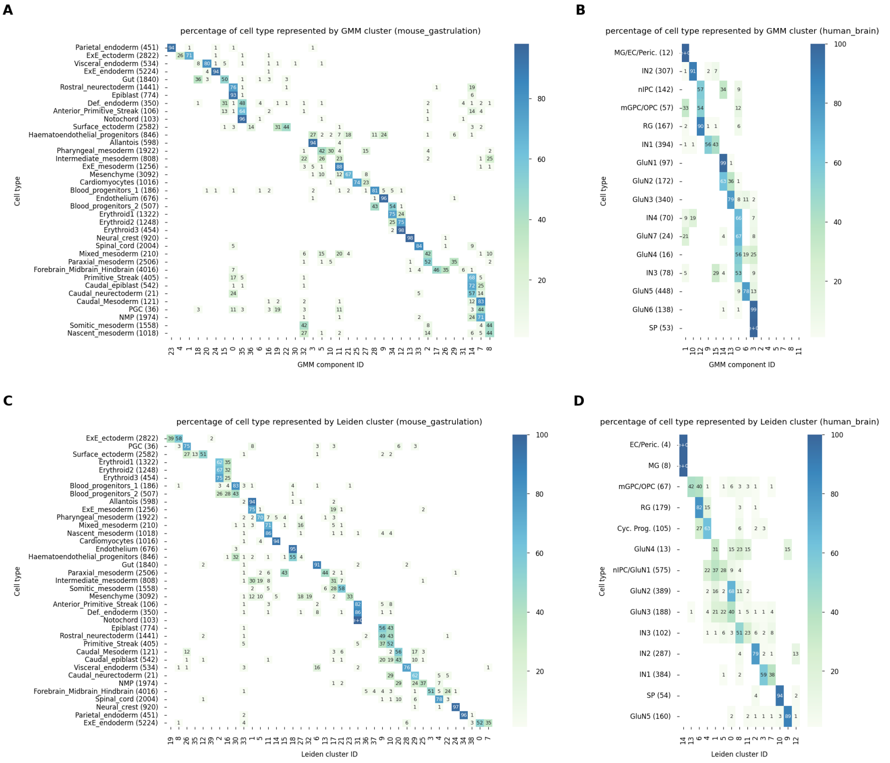


**Supplementary Figure 13 Clustering of components as percentage of cells in a cell type. A-B)** multiDGD. **A)** Latent clustering heatmap of the mouse gastrulation data set. **B)** Latent clustering heatmap of the human brain data set. The x axis presents the component IDs, the y axis the annotated cell types. Numbers inside the heatmap indicate the percentage of samples in a cell type assigned to a given component. **C-D)** MultiVI. Same as A and B but with Leiden clustering performed on the representation space.

**Table 3 Run time comparison of multiDGD and MultiVI on mouse gastrulation data sets with different data dimensionalities.** Both models were trained on two versions of the mouse gastrulation data (as in feature efficiency Table 1). The first was the previously used set, where features had been selected for counts to be above zero for at least five percent of cells. The second version was the full data set. The run time is taken from training multiDGD instances for 600 epochs and MultiVI for 500 epochs on the same cluster with equivalent resources. Run times, however, are not a robust metric of computational efficiency and are especially difficult to compare if not the same hardware was used. In less extreme cases, VAE-based models and DGDs have been shown to be comparable when it comes to run times [18].

| Model | Feature set | Run time |
|---|---|---|
| multiDGD | 5% | 10h 22min |
| MultiVI | 5% | 26h 54min |
| multiDGD | all | 11h 28min |
| MultiVI | all | 24h 38min |

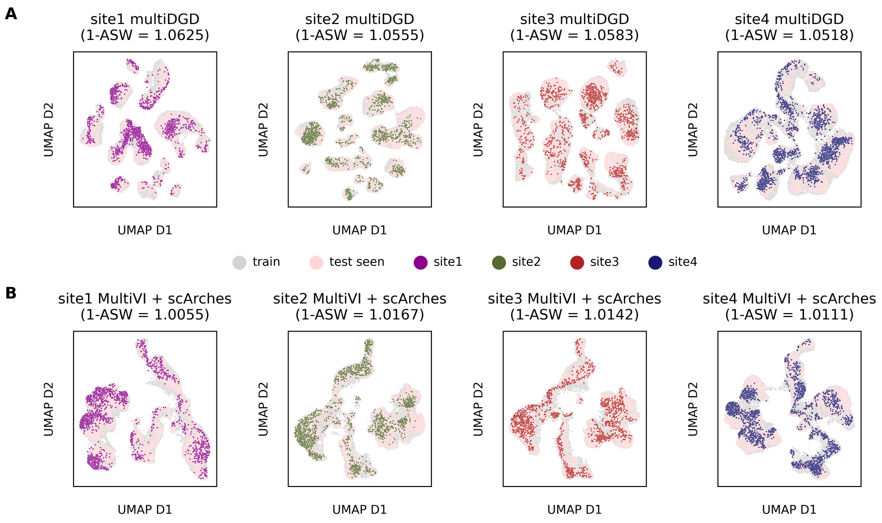


**Supplementary Figure 14 UMAP visualizations of latent spaces derived from models trained on three out of four batches.** Seen and unseen test representations are transformed onto the latent space and highlighted in color. Unseen test samples are colored by their respective 'site' (batch) and indicated in the plot titles. **A)** Latent spaces derived from multiDGD (trained on three batches). **B)** Latent spaces derived from MultiVI (trained on three batches) with scArches fine tuning.

**Table 4 Relative prediction performances on the held out bone marrow test set.** Relative prediction performances were calculated as the average prediction performance divided by the reconstruction performance over cells. We report means and standard deviations as the prediction-reconstruction ratio for each modality metric. BA stands for balanced accuracy.

| Model | prediction-reconstruction ratio | Metric |
|---|---|---|
| MultiVI | $1.0533 \pm 0.1404$ | RMSE (RNA) |
| multiDGD | $1.2272 \pm 0.7873$ | RMSE (RNA) |
| MultiVI | $0.9994 \pm 0.0029$ | BA (ATAC) |
| multiDGD | $0.9979 \pm 0.0138$ | BA (ATAC) |

**Table 5 Prediction and reconstruction performances on bone marrow data.** We report means and standard error of the means for both reconstruction and prediction performances of each modality metric. BA stands for balanced accuracy. Significant differences are highlighted by bold font of the values of the better performing model.

| Model | Prediction | Reconstruction | Metric |
|---|---|---|---|
| MultiVI | $\mathbf{0.6512 \pm 0.0136}$ | $0.5911 \pm 0.0107$ | RMSE (RNA) |
| multiDGD | $0.7262 \pm 0.0166$ | $\mathbf{0.5323 \pm 0.0081}$ | RMSE (RNA) |
| MultiVI | $0.5021 \pm 0.0001$ | $0.5024 \pm 0.0001$ | BA (ATAC) |
| multiDGD | $\mathbf{0.7092 \pm 0.0012}$ | $\mathbf{0.7107 \pm 0.0012}$ | BA (ATAC) |

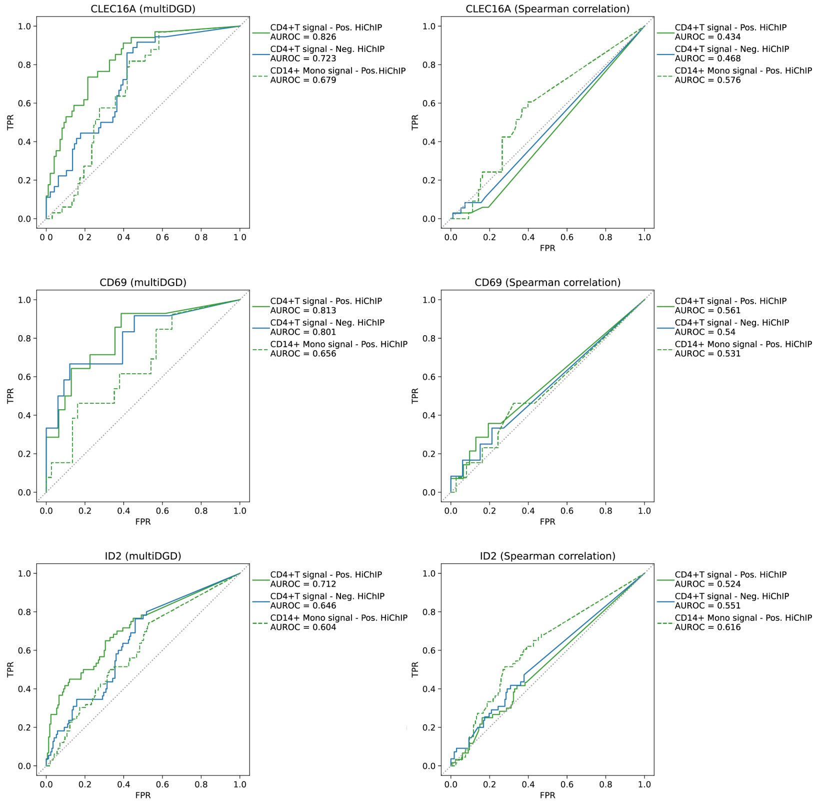


**Supplementary Figure 15 Prediction of enhancer-gene links from HiChIP data.** Receiver Operating Characteristic (ROC) curves for prediction of HiChIP enhancers with multiDGD (left column) or Spearmann correlation between peak accessibility and gene expression (right column), for 3 gene loci (rows).

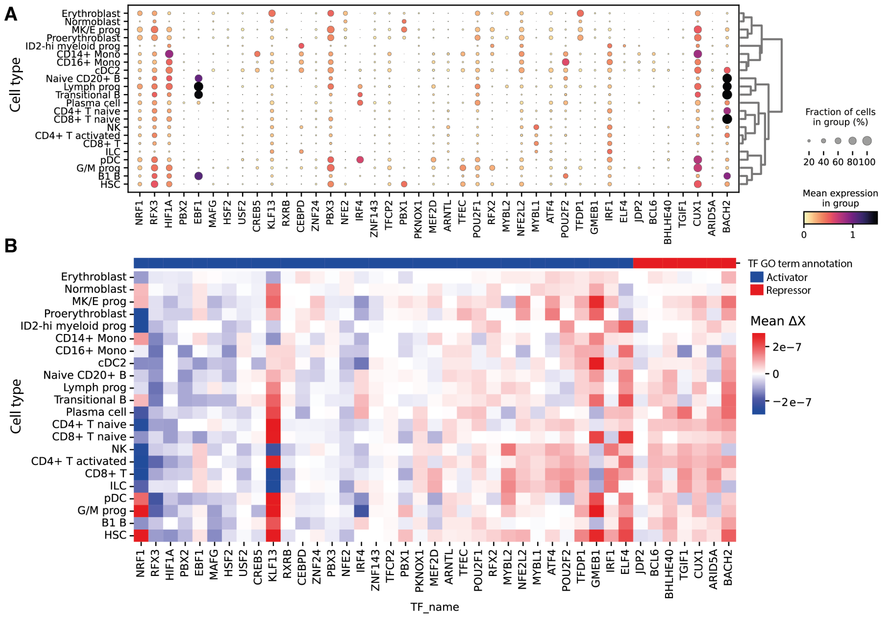


**Supplementary Figure 16 Transcription factor (TF) silencing experiment. A)** Dotplot for expression of perturbed TFs across annotated cell types in bone marrow data set. The dot color denotes the average log-normalized expression for the TF. The dot size indicates the fraction of cells expressing the TF in a cell type. **B)** Mean perturbation effects on peaks containing TF binding motifs across cell types in bone marrow data set. Cell types are grouped by hierarchical clustering as in (A). In both plots, TFs are ordered by mean effect on peaks with TF binding sites and annotated mode of action, as in Figure 5F.
